# β-actin mRNA interactome mapping by proximity biotinylation

**DOI:** 10.1101/405589

**Authors:** Joyita Mukherjee, Orit Hermesh, Carolina Eliscovich, Nicolas Nalpas, Mirita Franz-Wachtel, Boris Maček, Ralf-Peter Jansen

**Affiliations:** Interfaculty Institute of Biochemistry, University of Tübingen, Tübingen, Germany; Department of Medicine, Albert Einstein College of Medicine, Bronx, NY; Proteome Center Tübingen, University of Tübingen, Tübingen, Germany

**Keywords:** RNA-BioID, mRNA localization, FUBP3, RNA-binding protein, protein−RNA interaction, polarization index, super registration microscopy, label free quantification

## Abstract

The molecular function and fate of mRNAs are controlled by RNA-binding proteins (RBPs). However, identification of the interacting proteome of a specific mRNA in vivo is still very challenging. Based on the widely-used RNA tagging with MS2 aptamers for RNA visualization, we developed a novel RNA proximity biotinylation (RNA-BioID) method by tethering biotin ligase (BirA*) via MS2 coat protein (MCP) at the 3’-UTR of endogenous MS2 tagged β-actin mRNA (MBS) in mouse embryonic fibroblasts (MEFs). We demonstrate the dynamics of the β-actin mRNA interactome by characterizing its changes upon serum-induced localization of the mRNA. Apart from the previously known interactors, we identified over 60 additional β-actin associated RBPs by RNA-BioID. Among them the KH-domain containing protein FUBP3/MARTA2 has shown to be required for β-actin mRNA localization. We found that FUBP3 binds to the 3’-UTR of β-actin mRNA, is essential for β-actin mRNA localization but does not interact with the characterized β-actin zipcode element. RNA-BioID provides a tool to identify new mRNA interactors and to study the dynamic view of the interacting proteome of endogenous mRNAs in space and time.

**Significance statement:** Transport of specific mRNAs to defined sites in the cytoplasm allows local protein production and contributes to cell polarity, embryogenesis, and neuronal function. These localized mRNAs contain signals (zipcodes) that help directing them to their destination site. Zipcodes are recognized by RNA-binding proteins that, with the help of molecular motor proteins and supplementary factors, mediate mRNA trafficking. To identify all proteins assembling with a localized mRNA we advanced a proximity labeling method (BioID) by tethering a biotin ligase to the 3’ untranslated region of mRNA encoding the conserved beta-actin protein. We demonstrate that this method allows the identification of novel, functionally important proteins that are required for mRNA localization.

## Introduction

The spatial distribution of mRNAs contributes to the compartmentalized organization of the cell and is required for maintaining cellular asymmetry, proper embryonic development and neuronal function (1). Localized mRNAs contain cis-acting sequences, termed zipcodes or localization elements that constitute binding sites for RNA-binding proteins (RBPs) (1). Together with these RBPs, localized mRNAs form transport complexes containing molecular motors such as kinesin, dynein or myosin (2, 3). These ribonucleoprotein complexes (RNPs) usually include accessory factors such as helicases, translational repressors, RNA stability factors or ribosomal proteins (3). Thus, mRNPs as functional units do not only contain the information for an encoded polypeptide but also determine the precise spatio-temporal regulation of its translation and stability, thereby facilitating the correct subcellular localization of the translation product (4).

One of the best-studied localized mRNAs is β-actin that encodes the β isoform of the cytoskeleton protein actin (5). β-actin mRNA is localized to the protrusion of migrating fibroblasts (6) where its local translation critically contributes to the migrating behavior of this cell type (7–11) In the developing mouse (12) and Xenopus (13, 14) neurons, β-actin mRNA is transported to the growth cone during axonal extension and its deposition and local translation is highly regulated by external cues. In addition, translation of this mRNA in dendritic spines is involved in re-shaping the postsynaptic site of synapses (14). A well-defined localization element is present in the proximal region of the β-actin 3’-untranslated region (3’-UTR) (15). This cis-acting element is recognized and bound by the zipcode-binding protein, ZBP1(16), the founding member of the conserved VICKZ RBP family (17). ZBP1 (also called IGF2BP1 or IMP1) interacts with the β-actin zipcode via the third and fourth KH (hnRNP K homology) domains (16) and is required for RNA localization in fibroblasts and neurons (18). It has also been suggested that IGF2BP1 controls translation of β-actin mRNA by blocking the assembly of ribosomes at the start codon (11). IGF2BP1 appears to act as key RBP in β-actin mRNA distribution but other proteins, including IGF2BP2 (19), RACK1 (20), KHSRP/FUBP2 (21), KHDBRS1/SAM68 (22), FMR1 (23), and HuR (24) have been suggested to be involved in β-actin mRNA localization, although their molecular function is less clear.

To fully understand mechanism(s) of mRNA localization, it is important to identify and study the mRNA binding factors. Major technological advances like CLIP (crosslinking and immunoprecipitation) combined with next-generation sequencing allowed the identification of RNAs bound to specific RBPs (25) or the system-wide identification of RBPs that bind to polyA RNA (26, 27). However, the major approaches to determine which proteins associate with a specific RNA are affinity purification of modified or tagged RNAs together with their bound proteins, or co-immunoprecipitation of RNP components with the help of known RBPs (28). In addition, affinity capturing of specific RNPs with hybridizing antisense probes or via integrated aptamers has been successfully used (29–31). A limitation of these techniques is the potential loss of low affinity binders during purification, which has so far been addressed by in vivo UV cross-linking prior to cell lysis (25, 26). However, cross-linking enhances only the recovery of RBPs directly contacting nucleobases and therefore does not overcome the loss of other physiologically important RNA interactors (e.g., motor or adapter proteins). These limitations could be overcome by in vivo labelling of proteins while they are associated with the target RNA. BioID (proximity-dependent biotin identification) (32 – 34) has been successfully used to detect subunits of large or dynamic protein complexes like the nuclear pore complex (32) or centrosome (34). In BioID, a protein of interest is fused to a mutant version of the *E. coli* biotin ligase BirA (BirA*) that generates AMP-biotin (‘activated biotin’), which reacts with accessible lysine residues in its vicinity (33). After cell lysis, biotinylated proteins can be isolated via streptavidin affinity purification and identified using standard mass spectrometry techniques. Recently, BioID has also been applied to identify proteins associated with the genomic RNA of ZIKA virus (35).

In this study, we used BioID to characterize the proteome of the endogenous β-actin mRNPs. We showed that tethering of BirA* to an endogenous transcript does not only allow the identification of its associated proteins but can also be used to probe the environment of this mRNA. We identified FUBP3/MARTA2, an RBP from the conserved FUBP family of proteins (36–38), which was previously shown to mediate dendritic targeting of MAP2 mRNA in neurons (39, 40). We found that FUBP3 bind to and facilitate localization of β-actin mRNA to fibroblast leading edge. FUBP3 did not bind to the zipcode or IGF2BP1 but mediated β-actin RNA localization by binding to a distal site in its 3’-UTR. Therefore, RNA-BioID approach allows, with high confidence, to identify novel functional mRNA interactors within the cell.

## Results

### Tethering biotin ligases to the 3’-UTR of β-actin mRNA

In order to tether BirA* to the 3’-UTR of β-actin mRNA (Fig. 1*A*), we stably expressed a fusion of the nuclear localized signal (NLS) MS2 coat protein (MCP) (41), GFP and BirA* (MCP-GFP-BirA*) in immortalized mouse embryonic fibroblasts (MEFs) from transgenic β-actin-24 MBS mice (Fig. 1A, right panel) (8). These mice have both β-actin gene copies replaced by β-actin with 24 MS2 binding sites (MBS) in their distal 3’-UTR. In parallel, NLS-MCP-GFP-BirA* was stably expressed in WT (wildtype) MEFs with untagged β-actin mRNA, to generate a control cell line to eliminate background biotinylation due to the presence of constitutive expression of BirA* (Fig. 1A, second left panel). Both constructs contain two copies of the MCP protein leading to a maximum of 12 GFP and 12 BirA* that can potentially bind to an mRNA. Since biotinylation or the expression of the MCP-GFP-BirA* might affect localization of the β-actin mRNA, we checked for the proper targeting of β-actin mRNA to the leading edge of the cell by single molecule fluorescent in situ hybridization (smFISH) (42) and analyzed RNA localization by polarization index calculation (9) (Fig. 1*B, C* and S1 Appendix fig. S1*A*-*F*). The distribution of mRNAs within cells was assessed using probes against the β-actin ORF (for primary and immortalized MEFs) and β-actin MBS (for the genetically modified immortalized MEFs: β-actin MBS, or β-actin MBS IGF2BP1 KO (10). In order to account for random distribution of an mRNA within the cell, we used probes against Gapdh as a control. Gapdh mRNA is a highly abundant and uniformly distributed mRNA to induce β-actin mRNA localization, cells were serum starved for 24 hrs. followed by stimulation with serum addition for 1 hr. The median of the polarization index of β-actin mRNA distribution was significantly lower in immortalized (WT) or genetically modified immortalized MEFs compared to primary MEFs (Fig. 1*C*). Stimulation of polarization by serum was observed for all the cell types used in a similar manner (Fig. 1*C*, gray bars). Also, as shown before (10) knockout of IGF2BP1 reduces significantly β-actin MBS mRNA polarization (Fig.1C). We observed that β-actin, Igf2bp1 mRNA together with GFP protein, levels were not affected (Fig. 1*D*, S1 Appendix fig. S2*A, B*). Altogether these results suggest that biotinylation and/or the expression of the MCP-GFP-BirA* does not affect regulation of β-actin mRNA in MEFs. Furthermore, cells with similar expression levels of MCP-GFP-BirA* were sorted by FACS (fluorescence activated cell sorting). As shown before (43), we also found no differences in the biotinylation efficiency in at labelling conditions of 50 µM to 300 µM of biotin for 6 - 48 hrs. For optimal biotinylation, we decided to perform proximity labelling by addition of 50 µM biotin to the medium for 24 hrs. To test if proximity labelling can identify known β-actin mRNA-associated proteins, we affinity purified biotinylated proteins followed by western blot detection of IGF2BP1 (mouse ZBP1). IGF2BP1 was biotinylated in MEFs expressing β-actin-MBS/GFP-BirA* but not in those expressing only GFP-BirA* (Fig. 1*E*), which demonstrates that our tool can successfully biotinylate zipcode-interacting proteins. To differentiate between endogenously biotinylated proteins and RNA-dependently biotinylated proteins, we performed streptavidin pulldown in cells expressing β-actin-MBS / GFP-BirA* and in cells expressing only MCP-GFP and observed biotinylation of numerous additional proteins (see S1 Appendix, fig. S3). We expected that MCP-GFP-BirA* represents a major fraction of these biotinylated proteins and therefore aimed at depleting the fusion protein from the lysate by GFP pulldown prior to streptavidin affinity purification. To our surprise, most of the biotinylated proteins were enriched in the GFP pulldown fraction (S1 Appendix fig. S3), which is likely due to co-purification of MCP-GFP-BirA*, β-actin mRNA and biotinylated proteins via binding to the mRNA or the fusion protein. RNA degradation with RNase A (S1 Appendix fig. S4) shifted a large part of the biotinylated proteins into the streptavidin fraction (S1 Appendix fig. S3), supporting the idea that most of the biotinylated proteins are associated with β-actin mRNA. Additional treatment with high salt and 0.5% SDS further optimized the streptavidin affinity purification and decreased the background binding of the magnetic beads used in this purification (S1 Appendix fig. S3).

**Figure 1:**
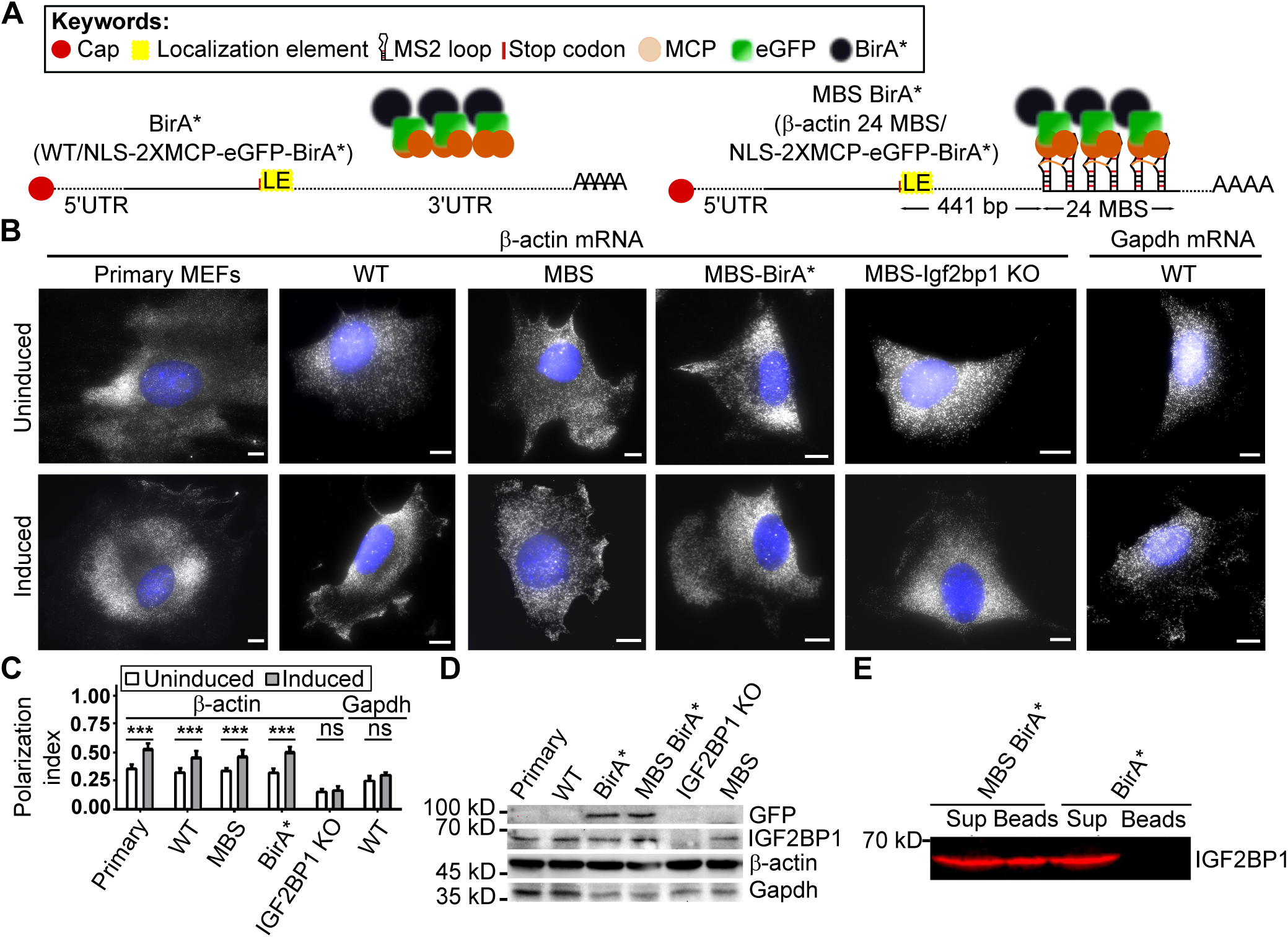
RNA BioID to detect proteins interacting with localized β-actin RNA. (A) Schematic of the β-actin-MBS/GFP-BirA*. Left: Control construct (named ‘BirA*’) used to detect background biotinylation due to overexpression of the NLS-MCP-GFP-BirA* construct. Control cells only expressing NLS-2xMCP-eGFP-BirA* lack the MBS cassette in the β-actin mRNA. Right: Construct used to detect β-actin mRNA associated proteins (named ‘β-actin MBS BirA*’). A 24xMS2 aptamer array (24MBS) was integrated in the 3’UTR of the endogenous β-actin gene 441 bp downstream of the stop codon. BirA* is targeted to 24MBS by its fusion to a MS2 coat protein dimer (2xMCP). (B) Representative β-actin smFISH images of (from left to right) primary MEFs, immortalized MEFs (WT), β-actin MBS, β-actin MBS BirA*, and β-actin MBS Igf2bp1 KO MEFs, as well as Gapdh smFISH images in immortalized (WT) MEFs (rightmost images). These and similar images were used to calculate the polarization index (C) of mRNA localization under serum uninduced (top panels) and serum uninduced (lower panels) conditions. β-actin mRNA was detected by probes against the β-actin ORF or MBS region, Gapdh mRNA was detected by probes against its ORF (gray). Scale bar: 10 µm (C) Bar graphs of polarization index for Gapdh mRNA and for β-actin mRNA in different MEFs (from left to right primary, immortalized (WT), β-actin MBS, β-actin MBS BirA*, β-actin MBS Igf2bp1 KO). Polarization index was calculated in total 100 of cells from 3 biological replicates. Line represents the median values. (D) Protein levels of endogenous β-ACTIN, IGF2BP1, and heterologous MCP-GFP-BirA* (detected by anti-GFP antibody). For the quantification for the western blot, see SI appendix, fig. S2. (E) Biotinylation of IGF2BP1 depends on MBS sites in β-actin. Following RNase A treatment, biotinylated proteins were affinity-purified with streptavidin-coated beads from cells expressing 2xMCP-eGFP-BirA* in the presence (β-actin-24MBS) or absence (β-actin) of MCP binding sites (MBS). Presence of IGF2BP1 was probed by a specific antibody in bead fraction (Beads) and supernatant (Sup).

### β-actin mRNA interactors under serum-induced and uninduced conditions

β-actin mRNA localization to the lamellipodia of chicken and mouse fibroblasts increases after serum induction (6, 44). It was also shown that during serum starvation, cells enter a quiescent phase of the cell cycle (6), with an overall reduction in actin stress fibers or focal adhesions (44). Since efficient biotinylation requires at least 6 hrs. of incubation with biotin, we next applied smFISH to verify that β-actin mRNA localization persists during our labelling period. As shown before (5), mouse fibroblasts induce β-actin mRNA localization after serum addition (Fig. 1*B*,1*C*) and the fraction of MEFs with β-actin localized to lamellipodia increases within one hour but remains constant over the next 6 hours.

To determine and compare the β-actin associated proteomes in uninduced and serum-induced MEFs, we performed RNA-BioID under both conditions (three replicate experiments each). Unspecific as well as endogenous biotinylation was assessed by performing BioID in MEFs expressing MCP-GFP-BirA* in the absence of MS2 aptamers in β-actin mRNA. Affinity-captured biotinylated proteins were identified and quantified by mass spectrometry using label free quantification (LFQ). Principal component analysis of the datasets revealed that the different conditions cluster apart from each other in dimensions 1 and 2 (explaining 33.8% and 15.5% of variance), while the replicates with the same conditions cluster together demonstrating biological reproducibility (S1 Appendix fig. S5). Calculating the Spearman correlation between all sample types and replicates (S1 Appendix fig. S6) supports the high reproducibility between biological replicates (correlation ≥ 0.97). In addition, it showed better correlation between uninduced and induced samples (average 0.95) compared to control. In total, we found 169 (or 156) significantly enriched proteins in induced (or uninduced) MEFs compared to control cells (see S1 Appendix, fig. S7 and S8*A*). Of these, 47 were enriched only under induced conditions (Supplementary Table 5). To assess the differential enrichment of the proteins under each condition, a Tukey post-hoc test was performed after the ANOVA, and the significance was set to an adjusted p-value of 0.05 following Benjamini-Hochberg multiple correction testing (see Materials and Methods). A large fraction of the enriched proteins (30% and 34%) under induced or uninduced conditions over control represent RNA-binding proteins (Fig. 2, red solid circles). Among these are RBPs (IGF2BP1, IGF2BP2, KHSRP, KHDRBS1, FMR1, HuR, RACK1, named in red) already known to control specific aspects of β-actin mRNA physiology. Other enriched RBPs have been associated with the localization of mRNAs in other cell types or organisms, including STAU1 and STAU2 (45–47), SYNCRIP (48), and FUBP3 (38). Furthermore, 85 proteins were significantly enriched under serum-induced compared to uninduced conditions (see S1 Appendix, fig. S8). However, the majority of the above-mentioned RBPs (including IGF2BP1) become biotinylated under induced as well as uninduced conditions, indicating that they are associated with β-actin mRNA under both conditions (Fig. 2*C*).

**Figure 2:**
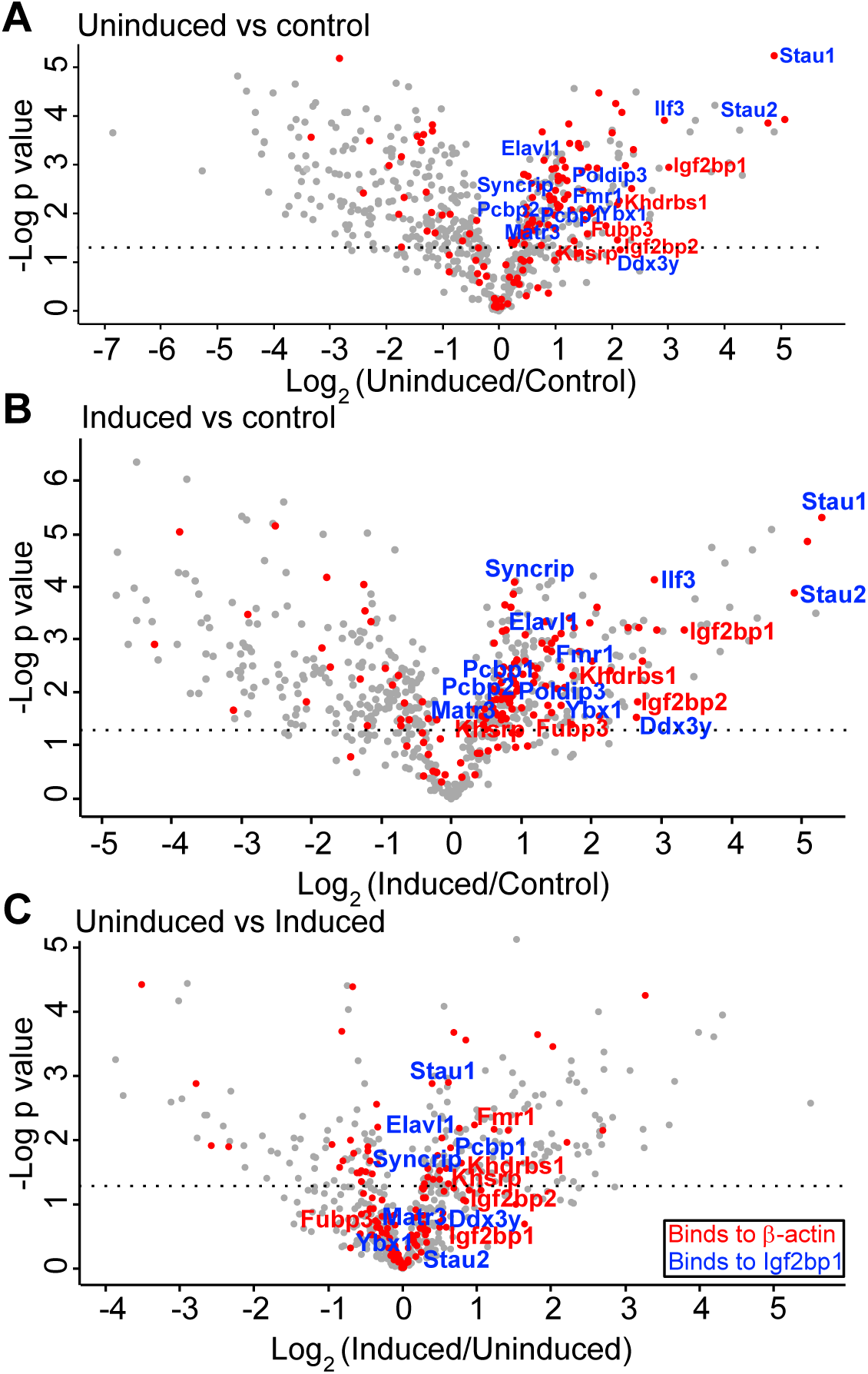
Enrichment of biotinylated proteins in control MEFs, or MEFs expressing β actin-MBS-BirA* under serum-induced or uninduced conditions. Volcano plot representation of biotinylated proteins in (A) uninduced MEFs compared to control MEFs, (B) serum-induced MEFs compared to control MEFs. (C) serum-induced MEFs compared to uninduced MEFs. In the volcano plots, the X-axis represents log2 fold change in protein abundance and the Y-axis represents the –log10 p-value. Red circles are known RBPs identified by GO-molecular function analysis. Proteins names in red represent known β-actin mRNA interactors and proteins named in blue are RBPs known to bind to IGF2BP1. Dotted line indicates p = 0.05.

A cluster analysis (Fig. 3) reveals at least five different patterns of biotinylated proteins in induced, non-induced and control MEFs (Fig. 3*B, C*). In control MEFs, we see enrichment of mainly nuclear proteins (cluster 1). This is expected since the unbound MCP-GFP-BirA* is enriched in the nucleus due to an N terminal nuclear localization sequence (8). Cluster 1 also contains abundant cytoplasmic proteins like glycerol aldehyde phosphate dehydrogenase (GAPDH). Cluster 3 represents proteins that are equally found in MEFs under all conditions and contains e.g. ribosomal proteins. Proteins allocated to the other three clusters (clusters 2, 4, 5) are overrepresented in the biotinylated proteome of MEFs expressing β-actin-MBS/GFP-BirA*. Of specific interest are clusters 4 and 5. In cluster 4, with proteins that are more biotinylated under serum-induced conditions, we find RNA-binding proteins, among them FMR1 and KHSRP that have been reported to function in β-actin mRNA localization or bind to IGF2BP1. Another group of proteins that are enriched in this cluster are proteins of the actin cytoskeleton (e.g. Filamin B, Cofilin-1, Myh9, Tpm4, Plastin-3). Their enrichment likely reflects the deposition of the β-actin mRNA in the actin-rich cortical environment of the MEF’s leading edge. Finally, cluster 5 contains proteins found in β-actin-MBS MEFs under induced as well as non-induced conditions but not in control MEFs. This cluster shows an enrichment for proteins involved in mRNA-binding, RNP constituents or ribosomal proteins. Since this cluster contains the zipcode-binding protein IGF2BP1, we hypothesized that other proteins in this cluster, e.g. FUBP3 are likely candidates for β-actin mRNA regulatory factors.

**Figure 3:**
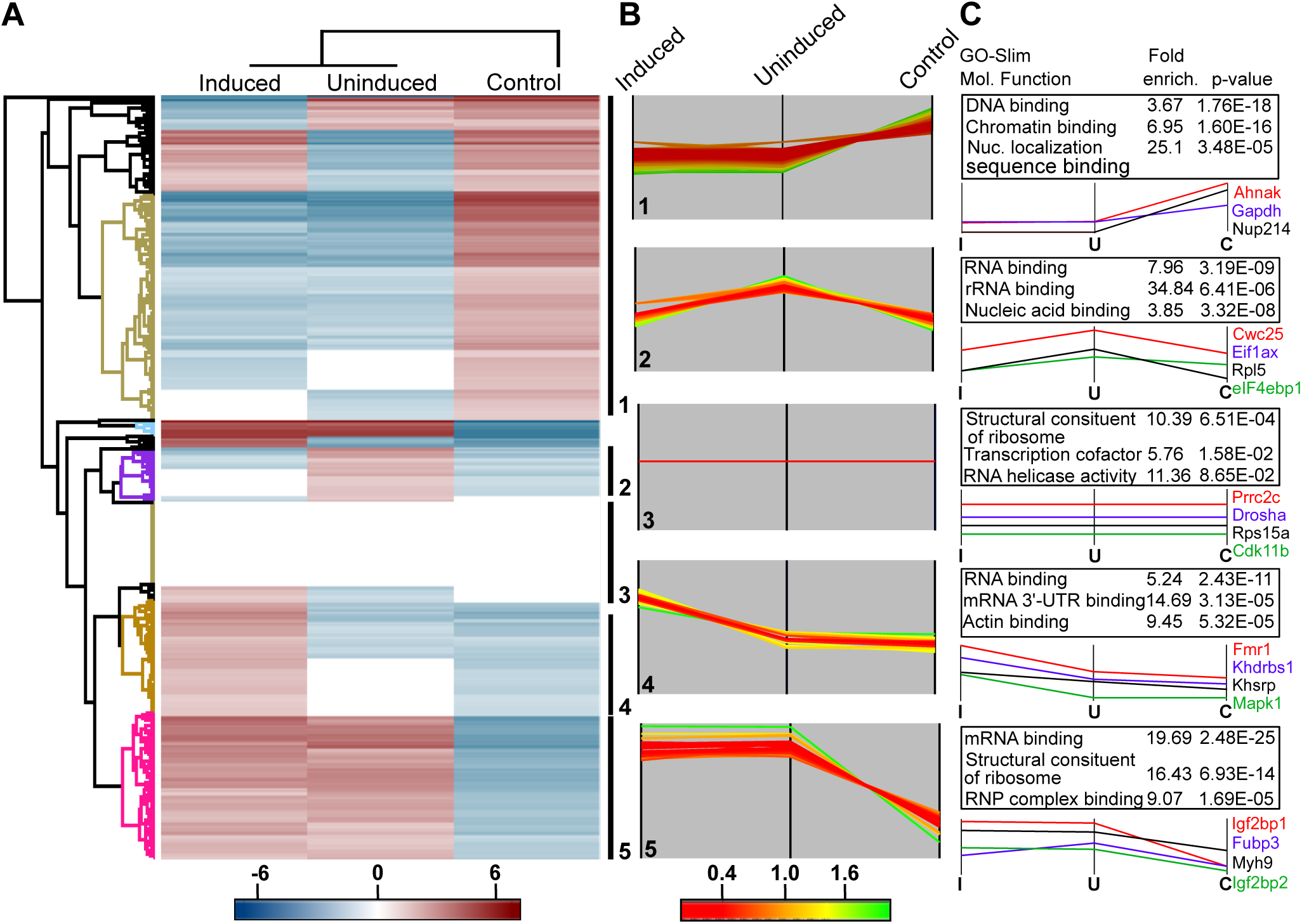
Cluster analysis of biotinylated proteins in control MEFs or MEFs expressing β-actin-MBS-BirA* under serum-induced or uninduced conditions. (A) Hierarchical clustering of biotinylated proteins in serum-induced, uninduced β-actin-MBS-BirA* MEFs and control MEFs (lacking β-actin-MBS). Enrichment is indicated in red coloring, depletion in blue. Various clusters of protein groups are highlighted in the dendrogram. (B) Profile plots of five selected clusters showing distinct enrichment patterns of biotinylated proteins: 1. strongly enriched in control MEFs; 2. enriched in β-actin-MBS-BirA* MEFs under uninduced condition; 3. similar enrichment in all MEFs; 4. enriched in β-actin-MBS-BirA* MEFs under serum-induced conditions; 5. enriched in β-actin-MBS-BirA* MEFs under serum induced and uninduced conditions compared to the control MEFs. Degree of enrichment in each specific cluster is represented by coloring (green, more enriched, red, and less enriched). (C) Functional analysis of protein annotation terms results in multiple categories that are enriched in the selected clusters. GO-slim molecular function terms, the corresponding enrichment factor, and the p-value are shown in the table. Selected examples of proteins found in each cluster are depicted below the tables.

### FUBP3 is a component of the β-actin mRNP

To confirm the association of the identified proteins and MS2-tagged β-actin mRNA, we combined single molecule FISH with immunofluorescence (smFISH-IF) using Cy3 labeled probes against either the ORF or the MBS of β-actin mRNA and antibodies against GFP, FUBP3 or IGF2BP1 in WT or β-actin-MBS MEFs (Fig. 4*A*-*C*). While ORF probes were used to detect β-actin mRNA in WT MEFs, MBS probes against the MS2 loop sequences were used to detect the β-actin mRNA in β-actin-MBS MEFs. The association between β-actin mRNA and the proteins was determined by super registration microscopy (47). Briefly, we corrected the images for chromatic aberration and mechanical shifts in Cy3 and Cy5 channels by using broad spectra fluorescent microsphere beads (S1 Appendix fig. S9); and we determined that colocalization of smFISH and immunofluorescence signals did not occur by chance accident within the cell by using a positive (MBS-GFP, Fig. 4*A*) and a negative (Gapdh-GFP, Fig. 4*D*) control for RNA-protein interaction. We calculated the association between the RNA and protein molecules as a function of their distances apart for positive and negative controls (Fig. 4*E*). For the positive control, 91% of the observed distances from the labeled probes to the MBS and from the antibodies to the GFP were within 60 nm (the optimal distance). In contrast, 10% of the observed associations in the negative control (using Gapdh probes and MCP-GFP) took place within the same distance (i.e., 60 nm) (Fig. 4*E, F*). When combining smFISH of Gapdh with immunofluorescence against MCP-GFP, few overlapping events were observed in less than 150 nm distance, when compared with MBS-GFP (Fig. 4*A*, 4*D* and 4*E*). At larger distances (>150 nm), the fluorescence signals in both channels are more likely to overlap by chance and therefore considered a random event. We found that at the optimal distance of 60 nm, β-actin mRNA was associated withIGF2BP1 and FUBP3 in MEFs. The RNA-protein associations were 37% and 29% for IGF2BP1 and FUBP3 with β-actin, respectively in MEFs (Fig. 4*B, C* and *F*). These associations were significantly higher than the nonspecific interaction between Gapdh and MCP-GFP (10%), suggesting the physical contact between the molecules.

**Figure 4:**
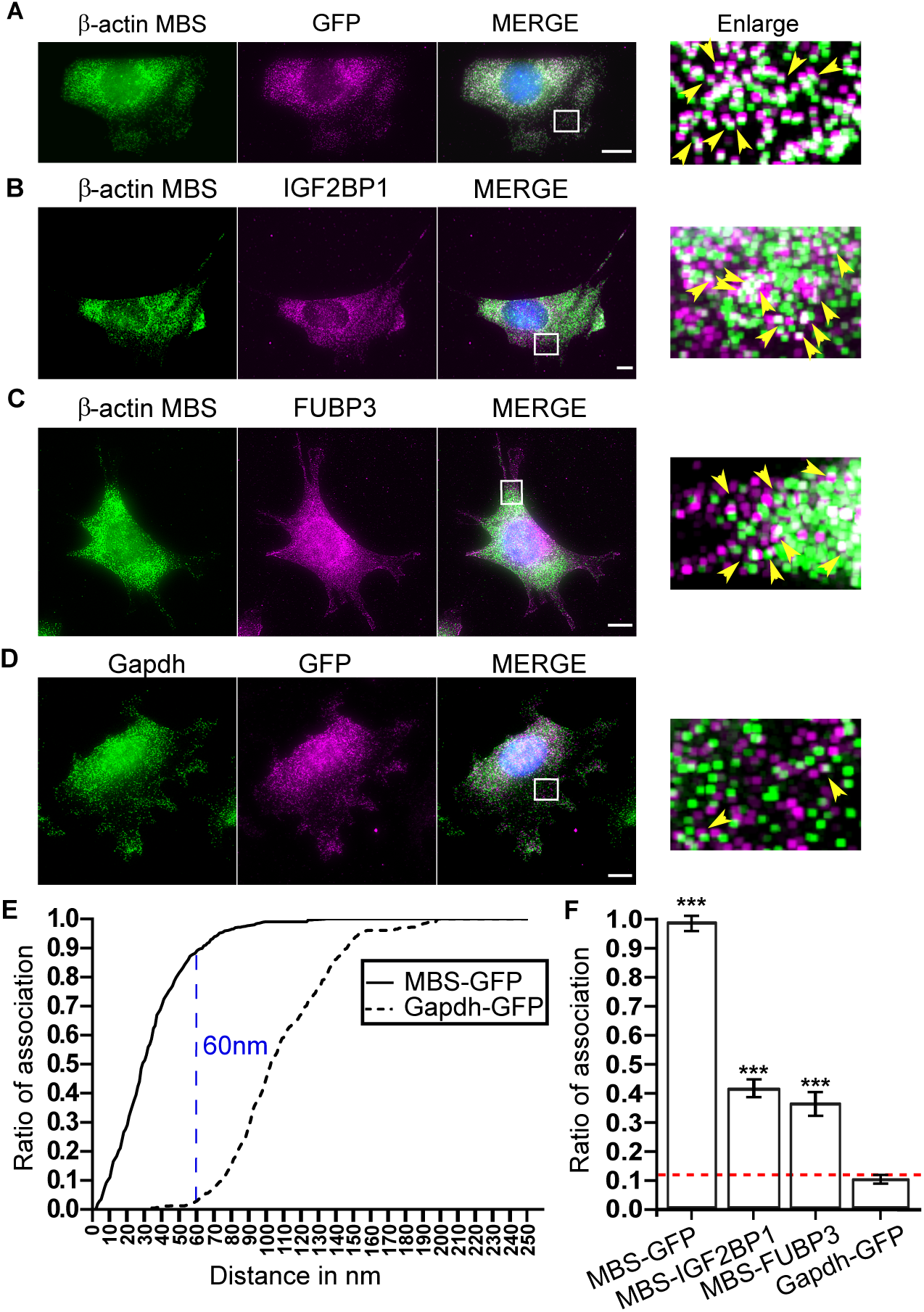
Association analysis of IGF2BP1 and FUBP3 with β-actin-MBS mRNA by super registration microscopy. (A-D) Representative smFISH-IF images of MEFs expressing β-actin MBS and MCP-GFP. Images A-C show MEFs stained for β-actin mRNA (MBS FISH probes, Cy3; green) and MCP-GFP (A), IGF2BP1 (B), FUBP3 (C). Immunofluorescence staining is shown in magenta. MCP-GFP was used as a positive control to determine the optimum distance between mRNA and protein. (D) represents staining of Gapdh mRNA (probes from Biosearch, Cy3; Green) together with MCP-GFP and was used as a negative control to determine the distance for association between two signals occurring by chance. On the right side of each panel a one pixel dilated, enlarged versions is been shown (47). (E) Association curves between an mRNA (black: β-actin MBS, dotted: Gapdh) and MCP-GFP protein. The curve of association is calculated as the cumulative ratio of association for intermolecular distances (in the range between 0 and 250 nm) that were less than a given observed distance as was described in (48). Blue line represents the distance where the mRNA–protein association for MCP-MBS and MCP-Gapdh is maximally separated (optimal distance, OD = 60 nm). (F)Summary of association analysis of β-actin mRNA and indicated proteins by smFISH-IF and super registration. Dotted red line indicates background association defined by MCP-Gapdh. Error bar represents SD. Unpaired t test was performed; P > 0.05. *P < 0.05; ****P < 0.0001.

### FUBP3 and IGF2BP1 binds on different regions of β-actin mRNA and interacts with each other in an RNA dependent manner

To validate the data demonstrating RNA-protein association by super registration microscopy, we performed co-immunoprecipitation of β-actin mRNA with FUBP3 and IGF2BP1 (Fig. 5*A*). Co-immunoprecipitation was tested with four mRNAs β-actin, Cofilin1, Igf2bp1, Fubp3 (Fig.5*A*). IGF2BP1 binds to all the mRNAs tested, which reflects previous observations in Hela cells, where almost 3% of the transcriptome was shown to bind to IGF2BP1 (49). Co-precipitation of β-actin with FUBP3 (23% of input bound to FUBP3) was similar to IGF2BP1 (37%). These numbers are consistent with the degree of RNA-protein association seen by co-localization (Fig. 4). In contrast, β-actin mRNA is not efficiently bound by the RBP VIGILIN, indicating that this mRNA does not associate with every RBP (S1 Appendix fig. S10). The localized Cof1 mRNA (50) is bound by both FUBP3 and IGF2BP1 to a similar extent (48%). To further substantiate our finding that FUBP3 can bind independently of IGF2BP1 to β-actin mRNA, we performed co-immunoprecipitation experiments of IGF2BP1 and FUBP3 (Fig. 5*B*) in presence or absence of RNase A. The RNA-binding protein STAU2 was used as positive control since it has been shown to bind to IGF2BP1 (51). Co-immunoprecipitation of IGF2BP1 and FUBP3 vanishes upon RNase treatment, indicating RNA dependent interaction between these two proteins. We conclude that FUBP3 does not bind to β-actin mRNA via IGF2BP1 but both proteins may rather they bind β-actin at independently at different sites. We next used recombinant histidine-tagged proteins (FUBP3-HIS and IGF2BP1-HIS) in pulldown assays (Fig. 5*C*, 5*D*) to test binding to in vitro transcribed RNA fragments of β-actin mRNA. We selected the complete 643 bp long β-actin 3’UTR and the 54-nucleotide localization zipcode element of β-actin. As negative control for IGF2BP1 binding we used a mutant version of the zipcode region (16). In addition, we used a 49-nucleotide long region adjacent to the zipcode (proximal zipcode; (15)). A 79 nt long region in the 3’UTR 460 nt downstream to the stop codon of β-actin mRNA, which spans a potential FUBP3 binding motif UAUG (52) and a 75 nucleotide fragment of the same region but carrying a deleted UAUG motif were used to specifically probe FUBP3 binding. The capturing assay was performed in total bacterial lysates to allow bacterial RBPs to compete for RNA binding. RNA captured by the His-tagged fusion proteins was detected by quantitative RT-PCR and normalized to the input (Fig. 5*D*). We found that IGF2BP1 and FUBP3 bind to the 3’UTR of β-actin mRNA, while neither can interact with the mutated zipcode or zipcode proximal region. Only FUBP3 binds to the 79-nucleotide long region containing UAUG motif on the 3’UTR of β-actin mRNA and the binding is abolished in absence of this motif (Fig. 5*D*). This is highly suggestive of direct binding of FUBP3 to the UAUG motif in the 3’-UTR of β-actin to identify the KH domain(s) of FUBP3 responsible for binding β-actin mRNA, we introduced mutations in the conserved KH domains of the protein. Each functionally important G-X-X-G motif in the four KH domains was changed to the inactive G-D-D-G (53) and individual mutant proteins were transiently expressed in MEFs as C-terminally tagged mCherry fusion protein. The G-D-D-G mutation in KH domain 2 resulted in loss of the cytoplasmic punctate signal seen in wild type FUBP3, which is reminiscent of punctate pattern observed for mRNPs (Fig. 5*E*). We conclude that KH2 in FUBP3 is important for its integration into RNP particles and likely constitutes the critical domain for RNA binding.

**Figure 5:**
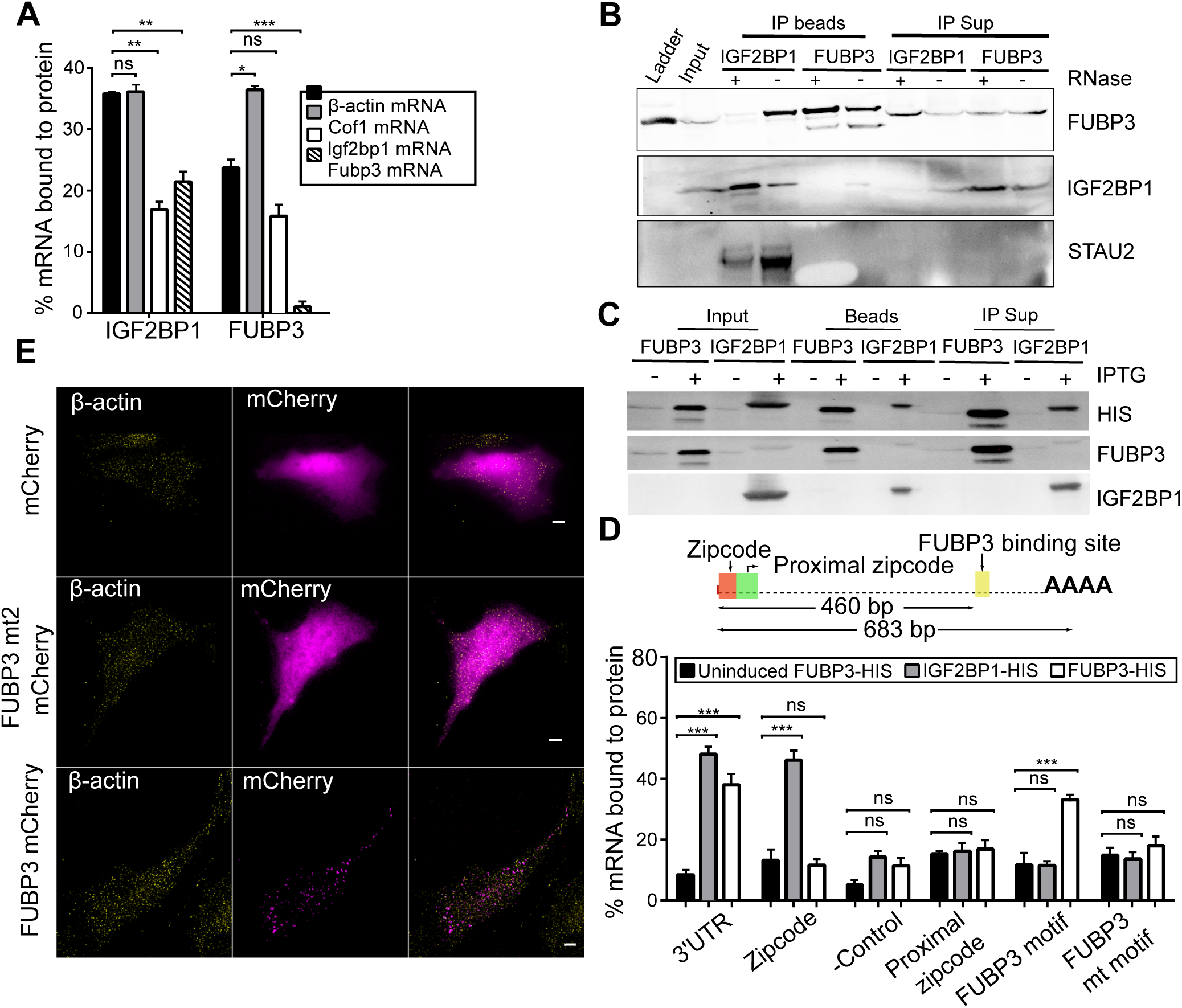
FUBP3 binds to β-actin 3’UTR. (A) Co-immunoprecipitation of selected mRNAs with IGF2BP1 and FUBP3. Bars represent percentage of input mRNA co-purifying with the indicated protein. IGF2BP1 binds to several endogenous mRNAs such as Cofilin1, Igf2bp1, and Fubp3. FUBP3 binds to 23% of endogenous β-actin mRNA while IGF2BP1 was associated with 37% of endogenous β-actin mRNA. Error bars represents mean ± sem from three independent experiments. (B) Co-immunoprecipitation of STAU2, FUBP3 and IGF2BP1. Immunoprecipitation was performed from wild type MEFs either with anti-FUBP3 or anti-IGF2BP1 antibodies in presence and absence of RNase A. IGF2BP1 co-precipitates with FUBP3 only in absence of RNase A while binding of STAU1 to IGF2BP1 is RNA-independent. (C) Pulldown of His-tagged fusion proteins of IGF2BP1 and FUBP3 from bacterial lysates of E. coli grown under IPTG induced or IPTG uninduced conditions. Magnetic beads were used to precipitate either IGF2BP1-HIS or FUBP3-HIS (D) Top: Schematic representation of 3’UTR of β-actin mRNA. The 683 bp long 3’UTR contains the 54 nt long zipcode sequence (after the stop codon), the proximal zipcode sequence (49 bp following the zipcode) and a potential FUBP3 binding sequence (460 bp downstream of the stop codon) with a consensus UAUG motif. Bottom: Binding of in vitro transcribed RNA fragments of β-actin (complete 3’UTR, zipcode, proximal zipcode, zipcode mutant, FUBP3 binding motif region, region with mutated FUBP3 binding motif) to IGF2BP1 or FUBP3. RNAs were added to E. coli lysates with or without (IPTG-uninduced) expressed His tagged fusion protein. After affinity purification, bound RNAs were detected by quantitative RT-PCR. Bars represent percentage of input RNA. In contrast to IGF2BP1, FUBP3 shows little affinity for the zipcode sequence but binds to the 3’UTR and a region containing the UAUG motif in the 3’-UTR. Error bars represents mean±sem from three independent experiments. Statistical significance of each dataset was determined by Student’s t-test; *P < 0.05; ***P < 0.001. (E) RNA-binding domain KH2 is required for FUBP3 cytoplasmic granule formation. The conserved G-X-X-G motif of FUBP3 KH domains were individually mutated into G-D-D-G and wildtype and mutant proteins expressed in MEFs as mCherry fusion. Live cell imaging shows that wild type FUBP3-mCherry forms cytoplasmic granules whereas a KH2 mutant (FUBP3 mt2) is evenly distributed in the cytoplasm like the control mCherry protein. Scale bar represent 5 μm.

### Loss of FUBP3 affects β-actin mRNA localization

To validate that proteins identified by RNA-BioID are functionally significant for the mRNA used as bait, we performed shRNA mediated knockdown experiments for FUBP3. The effectiveness of the knockdown was validated by quantitative RT-PCR and western blot (Fig. 6*A*-*C*) using GAPDH as control since it does not interact with β-actin mRNA as shown by RNA BioID (Fig. 3*C*). FUBP3 knockdown only mildly reduced mRNA levels of β-actin or IGF2BP1 mRNAs (Fig. 6*B*). Similarly, IGF2BP1 protein levels did not significantly change upon FUBP3 knockdown (Fig. 6*C*), ruling out an indirect effect of FUBP3 on β-actin mRNA by limiting IGF2BP1 levels. However, we observed a slight increase in β-ACTIN protein level, indicating that FUBP3 might co-regulate β-actin mRNA translation or β-ACTIN protein stability.

**Figure 6:**
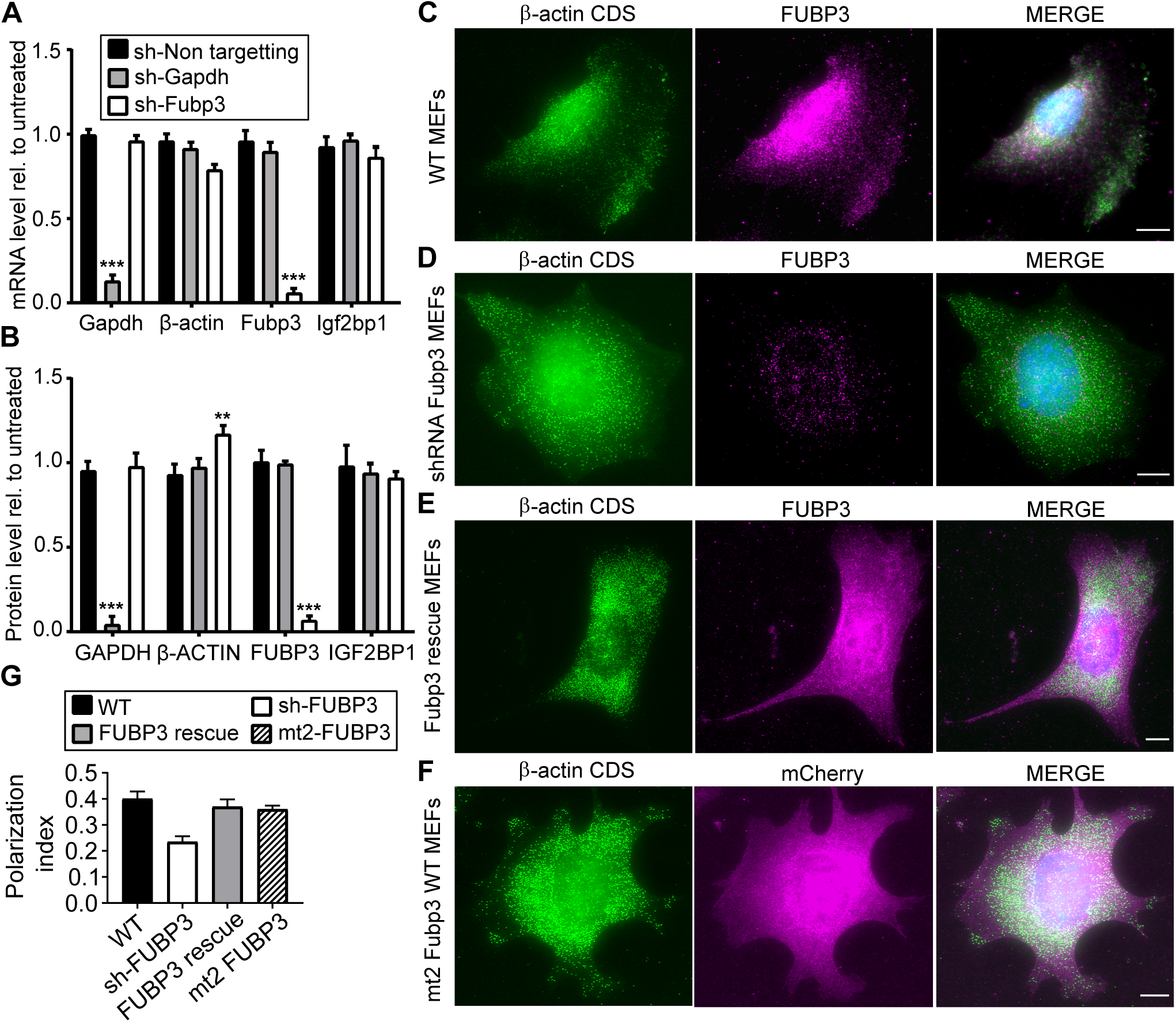
Downregulation of Fubp3 affects β-actin mRNA localization. (A) Western blot analysis to monitor shRNA-mediated knockdown of FUBP3 in non-targeted shRNA stably expressing in WT (2nd lane) compared to unmodified WT (lane 1) and WT stably expressing shRNA against Gapdh (lane 3) and stably expressing shRNA against FUBP3 (lane 4). Blot was reprobed against GAPDH and β-ACTIN and IGF2BP1 to access the changes in these proteins due to knockdown of FUBP3 or GAPDH. (B) Quantitative RT-PCR analysis of Gapdh, β-actin, Igf2bp1, and Fubp3 levels in knockdown cells. Corresponding mRNA levels in untreated cells were used for normalization. Statistical significance of each dataset was determined by Student’s t-test; *P < 0.05; ***P < 0.001. (C) Western blot quantification of GAPDH, β-ACTIN, IGF2BP1, and FUBP3 protein levels in knockdown cells. Protein levels in untreated cells were used as normalization control. Statistical significance of each dataset was determined by Student’s t-test; *P < 0.05; ***P < 0.001. (D-G) Representative smFISH-IF images of immortalized MEFs before or after Fubp3 knockdown. (D-F) β-actin mRNA (Cy3; green) and FUBP3 (magenta) are shown. (D) MEF before knockdown (WT), (E) MEF after shRNA treatment, (F) shRNA-treated MEF expressing a FUBP3 rescue construct (see methods) (H) Representative smFISH-IF image of MEF expressing the FUBP3 KH mutant mt2 (see Fig. 5). Scale bars represent 10 µm. (I) Polarization index for β-actin mRNA in MEFs from experiments shown in D-G. Polarization index was calculated from a total of 65 cells from 3 biological replicates. Bars represent the median values. Statistical significance of each dataset was determined by Student’s t-test; *P < 0.05; ***P < 0.001.

The effect of the FUBP3 knockdown or overexpression of mutant FUBP3 on β-actin mRNA localization was assessed by smFISH-IF (Fig. 6*D*-*G*, S1 Appendix fig. S11) and analyzed by calculating the polarization index (Fig. 6*H*). In control cells (immortalized MEFs, Fig. 6*D*), we found that FUBP3 and β-actin mRNA are expressed, and that the polarization index of β-actin mRNA was 0.37 (Fig. 6*H*). In FUBP3 knockdown cells (Fig. 6*E*), almost no FUBP3 signal was detectable and the β-actin mRNA polarization index dropped to 0.25 (Fig. 6H). To test if the reduction in polarization is due to loss of FUBP3, we expressed a knockdown-insensitive mCherry-tagged FUBP3 in these MEFs. Expression of this fusion protein was accessed by indirect immunofluorescence against mCherry whereas β-actin mRNA was visualized by smFISH (Fig. 6*F*). To determine the polarization index, we only used MEFs positive for mCherry. Although no full rescue was observed, the polarization index increased to 0.33 (Fig. 6*H*). This indicates that FUBP3 is important for β-actin mRNA localization. In addition, we analyzed the effect on β-actin mRNA distribution when overexpressing a mCherry-tagged FUBP3mt2 mutant lacking a functional KH2 domain (see above). As before, we selected MEFs with a mCherry signal for determination of the polarization index. We found that the polarization index was 0.31 (Fig. 6*H*) which is not significantly different from that of β-actin mRNA in wildtype MEFs. These data suggests that although KH2 is important for the forming FUBP3 containing RNP particle-like structures in the cytoplasm, it does not act as dominant negative mutation, probably since a mutant with this mutation does not compete with endogenous FUBP3.

## Discussion

Proximity biotinylation has facilitated the characterization of dynamic protein complexes by in vivo labelling of interaction partners. Here, we exploit this approach and demonstrate its utility for identifying functionally relevant RNA-binding proteins of a specific mRNA, mammalian β-actin. This is achieved by combining MS2 tagging of the mRNA of choice and co-expression of a fusion protein of the MS2 coat protein (MCP) and the biotin ligase (BirA*).

The primary goal for an RNA-based BioID is the identification of novel RNA interactors. As seen before in several proximity labelling (BioID or APEX-driven) approaches (43, 54, 55), the number of identified potential interactors for β-actin is far higher than the number of proteins identified in classical co-immunoprecipitation or co-affinity purification approaches. This might be due to the higher sensitivity of proximity labelling or its propensity to allow capturing of transient interactors (56). Although this can result in a skewed view of the actual components of a complex due to the rapid change in the composition of mRNP, it is beneficial to identify all the mRNP components during the life stages of an mRNA. The most highly represented class of proteins were RBPs (Fig. 3 and S1 Appendix fig. S8*B*), among them all RBPs that have been previously associated with localization, translational control or (de)stabilization of β-actin mRNA. Other RBPs like survival of motor neuron 1 (SMN1), which supports the association of IGF2BP1 with β-actin mRNA (57), were also found to be enriched in MEFs expressing β-actin-MBS compared to control MEFs, although with lower significance (p-value < 0.1).

We also analyzed our dataset for motor proteins involved in mRNA transport. Neither MYH10 (58) nor KIF11 (59) that have been suggested to work as β-actin mRNA transport motors were found as biotinylated proteins. The only motor we identified is MYH9, the heavy chain of a MYH10 related class II-A myosin although it was not significantly enriched (p = 0.08). The lack of motor proteins is compatible with a recent observation that β-actin localization in fibroblasts works primarily by diffusion to and trapping in the microfilament-rich cortex (60). This is also corroborated by our finding that components of the actin-rich cell protrusion (Fig. 3, cluster 4) are heavily biotinylated in MEFs after serum-induced localization of β-actin.

Overall, the cluster analysis shows that the majority of previously identified β-actin RBPs behave similarly under the two tested conditions (serum-induced and uninduced MEFs). This not only indicates that they interact with β-actin mRNA in MEFs even under steady state conditions, but It also makes it likely that other proteins, especially RBPs, found in this cluster might represent so far unknown β-actin mRNA interactors. By choosing the far-upstream binding protein FUBP3 as a potential candidate we demonstrate that this assumption holds true for at least this protein. FUBP3 not only binds to β-actin mRNA but its knockdown also results in a similar decrease of β-actin localization to the leading edge as seen for loss of IGF2BP1.

FUBP3, also named MARTA2 has been reported to bind to the 3’-UTR of the localized MAP2 mRNA in rat neurons (39) and regulates its dendritic targeting (40). Although the binding site of FUBP3 in MAP2 mRNA is not known, its preferred binding motif (UAUA/UAUG) was recently identified (52). This motif is present in the 3’-UTR of β-actin 460 nt downstream of the zipcode and a 79-nucleotide region containing this motif is bound by FUBP3. FUBP proteins might play a more substantial role in RNA localization since homologs of a second member of the FUBP family, FUBP2 were not only reported to be involved in MAP2 or β-actin mRNA localization but also present among the biotinylated proteins we identified. However, FUBP2 is mainly nuclear and its role in β-actin mRNA localization might be indirect (61). In contrast, FUBP3 seems to have a direct function in localizing β-actin as it binds to the 3’-UTR and its loss reduces β-actin mRNA localization independently of IGF2BP1. This independent function is supported by the observation that both proteins do not directly bind to each but to different regions of β-actin mRNA. A potential additional function could be translational regulation. Although less dramatic than seen for loss of IGF2BP1, knockdown of FUBP3 results in increased amounts of β-ACTIN protein while β-actin mRNA levels are similar or even lower than in untreated MEFs. This could be due to a loss of translational inhibition as it has been shown for IGF2BP1(11).

Its role in β-actin and MAP2 mRNA localization suggests that FUBP3/MARTA2 is a component of several localizing mRNPs. Of note, RNA-BioID on β-actin mRNA has identified even more RBPs that have been previously involved in the localization of other mRNAs, e.g. SYNCRIP (48) or Staufen (62). Several of these like STAU1 and STAU2 are highly enriched in our β-actin biotinylated proteome. This finding might on one hand reflect the participation of multiple RBPs in β-actin localization or regulation. It also shows that a common set of RBPs is used to control the fate of several different localized mRNAs in different cell types. Although RNA-BioID does not currently allow us to determine if all these RBPs are constituents of the same β-actin mRNP, belong to different states of an mRNP or to different populations, their identification allows addressing these questions to reach a more detailed understanding of the common function of RBPs on diverse mRNAs.

## Materials and methods

### RNA-BioID

For RNA-BioID, cells were incubated with 50 µM biotin at least for 6 hrs. Following incubation, cells were washed twice with 1x PBS and lysed in IP lysis buffer (50 mM Tris pH 7.5, 150 mM NaCl, 2.5 mM MgCl2, 1 mM DTT, 1% tween-20, and 1x protease inhibitor) and passed 10-12 times through a 21G needle. The lysate was cleared by centrifugation (12,000 x g for 10 min at 4°C) to remove cell debris. 10 µg of protein from the supernatant (total cell lysate) were used to check for protein biotinylation. In the remaining lysate, NaCl was added to a final concentration of 500 mM. 200 µl of a streptavidin magnetic bead suspension (GE Healthcare) were added and the high salt lysate incubated overnight at 4°C with end to end rotation. On the next day, the beads were collected (by keeping the beads on the magnetic stand for 2 min) and washed as described before (69). In detail, they were washed twice for 5 min with 0.3 ml wash buffer 1 (2% SDS), once with wash buffer 2 (0.1% (w/v) deoxycholate, 1% (w/v) tween-20, 350 mM NaCl, 1 mM EDTA pH 8.0), once with wash buffer 3 (0.5% (w/v) deoxycholate 0.5% (w/v) tween-20, 1 mM EDTA, 250 mM LiCl, 10 mM Tris-HCl pH 7.4) and 50 mM Tris-HCl pH 7.5, once with wash buffer 4 (50 mM NaCl and 50 mM Tris-HCl pH 7.4), and finally twice with 500 µl of 50 mM ammonium bicarbonate. 20 µl of the beads were used for western blot and silver staining, and 180 µl was subjected to mass spectrometry analysis. To release captured proteins for western blot analysis from streptavidin beads, the beads were incubated in 2x Laemmli buffer containing 2 mM saturated biotin and 20 mM DTT for 10 min at 95°C.

For biotinylation after serum induction, cells were starved for 24 hrs. as described in supplementary methods and induced with 10% serum containing media containing 50 µM biotin for at least for 6 hrs. to 24 hrs. Samples were processed for mass spectrometric analysis as described in supplementary material and methods.

### Microscopy and super registration microscopy

For live cell imaging, cells were imaged with a Zeiss Cell Observer wide field fluorescence microscope, operated by ZEN software (Zeiss), Illuminated with xenon arc lamp and detected with a CCD camera (Axiocam 506) with 100x/1.45 α-plan fluor oil immersion objectives (Zeiss). For the live cell imaging we used a dual band GFP/mCherry filter set (F56-319, AHF). For imaging of fixed cells, the microscope set up is the same as in Eliscovich et al.(47).

### Imaging Analysis

Single molecule localization was determined by FISH-QUANT (63) software; and super registration analysis was performed as described in Eliscovich et al.(47) with existing software packages and custom algorithm programs written in MATLAB (MathWorks). For polarization index calculation, after taking the max projections from all the z-stacks, polarization and dispersion index measurement were measured as mentioned before (9) with existing software package written in MATLAB (MathWorks).

### smFISH-IF

Immortalized WT MEFs or MEFs containing MS2 tagged β-actin but no MCP-GFP were seeded on a fibronectin coated cover glass in a 12 well cell culture plate and grown for 24 hrs. in serum free media, before serum containing media was added to the cells for 1-2 hrs. For smFISH and immunofluorescence the protocol was followed as previously described (47). In short, cells were washed three times with PBS, fixed for 10 min with 4% paraformaldehyde in PBS, washed three times in PBS then quenched in 50 mM glycine, and permeabilized with 0.1% Triton X-100 (28314; Thermo Scientific) and 0.5% Ultrapure BSA (AM2616; Life Technologies) in 1× PBS-M for 10 min. After washing with PBS, cells were exposed to 10% (vol/vol) formamide, 2× SSC, and 0.5% Ultrapure BSA in RNase-free water for 1hr at room temperature, before they were incubated for 3 h at 37 °C with either 10 ng custom labelled probes or 50 nm. Stellaris RNA FISH probes (Biosearch Technologies) (table S4). Primary antibodies against GFP (Aves Labs, Inc. GFP-1010), IGF2BP1 (MBL, RN001M) or FUBP3 (Abcam) were diluted (for dilutions refer to table S3) in hybridization buffer (10% formamide, 1 mg/mL Escherichia coli tRNA, 10% dextran sulfate, 20 mg/mL BSA, 2× SSC, 2 mM Vanadyl Ribonucleoside Complex, 10 U/mL Superasein (Ambion) in RNase-free water). After incubation and quick washing, cells were further incubated twice with an Alexa Fluor 647-conjugated secondary antibody (Life Technologies) in 10% formamide and 2× SSC in RNase-free water for 20 min at 37 °C. After four 2× SSC washes, DNA was counterstained with DAPI (0.1 μg/mL in 2× SSC; Sigma-Aldrich), and, after a final wash, cells were mounted using ProLong Diamond Antifade Reagent (Life Technologies).

## Supporting information

Combined Supplemental Information

## Data availability

Proteomic data supporting this study has been deposited into PRIDE, accession no: PXD010694.

Additional information is available in the SI Appendix and includes supplementary figures (S1-S12), supplementary tables (S1-S4), supplementary materials and methods, and a supplementary spreadsheet (supplementary table 5) with masspec results.

## Author Contributions

JM and RPJ conceived the project. JM performed experiments, analyzed the data and wrote the manuscript. JM performed the smFISH-IF and super registration experiments and analyzed the data. OH performed the live cell imaging experiment and wrote the manuscript. CE contributed to smFISH-IF, super registration experiments and imaging data analysis. JM, MFW, NN, and BM designed, performed and analyzed the mass spectrometry experiments RPJ supervised the project, interpreted the data and wrote the manuscript.

## Acknowledgement

We thank Jeff Chao (FMI, Basel), Imre Gaspar (EMBL, Heidelberg), Julién Bethune (BZH, Heidelberg), Dierk Niessing (University of Ulm), Michael Kiebler (University of Munich), Stefan Kindler (University of Hamburg), Stefan Hüttelmaier (University of Halle), and Ibrahim Muhammad Syed (IFIB Tübingen), for plasmids, cell lines, antibodies, or spike RNA. We are grateful to Robert H. Singer for hosting JM during an imaging internship. We are grateful to Frank Essmann, Ruth Schmid (both at IFIB Tübingen), and Silke Wahle (PCT Tübingen) for technical support, Jeetayu Biswas (Albert Einstein College, NY) for help with the polarization index script and IGF2BP1 KO cell line, and Matthew Cheng (IFIB Tübingen) for suggestions on the manuscript. The project was funded as project of the DFG Research Unit FOR2333 by a grant of the Deutsche Forschungsgemeinschaft (DFG JA696/11-1).

## Notes

#### Summary of Updates

Figures 1, 2, 4, 5, and 6 have been updated and include new experimental data. Large parts of the manuscript have been re-written. Supplementary information and figures have been re-arranged or replaced.

