## Supplementary material for "β-actin mRNA interactome mapping by proximity biotinylation": Combined Supplemental Information

**Supplementary Information (Mukherjee et al., 2019)**


| | Page no. |
|---|---|
| Supplementary Figures | 02 – 09 |
| Supplementary Tables | 12 – 17 |
| Supplementary Methods | 18 – 21 |

### Supplementary Figures

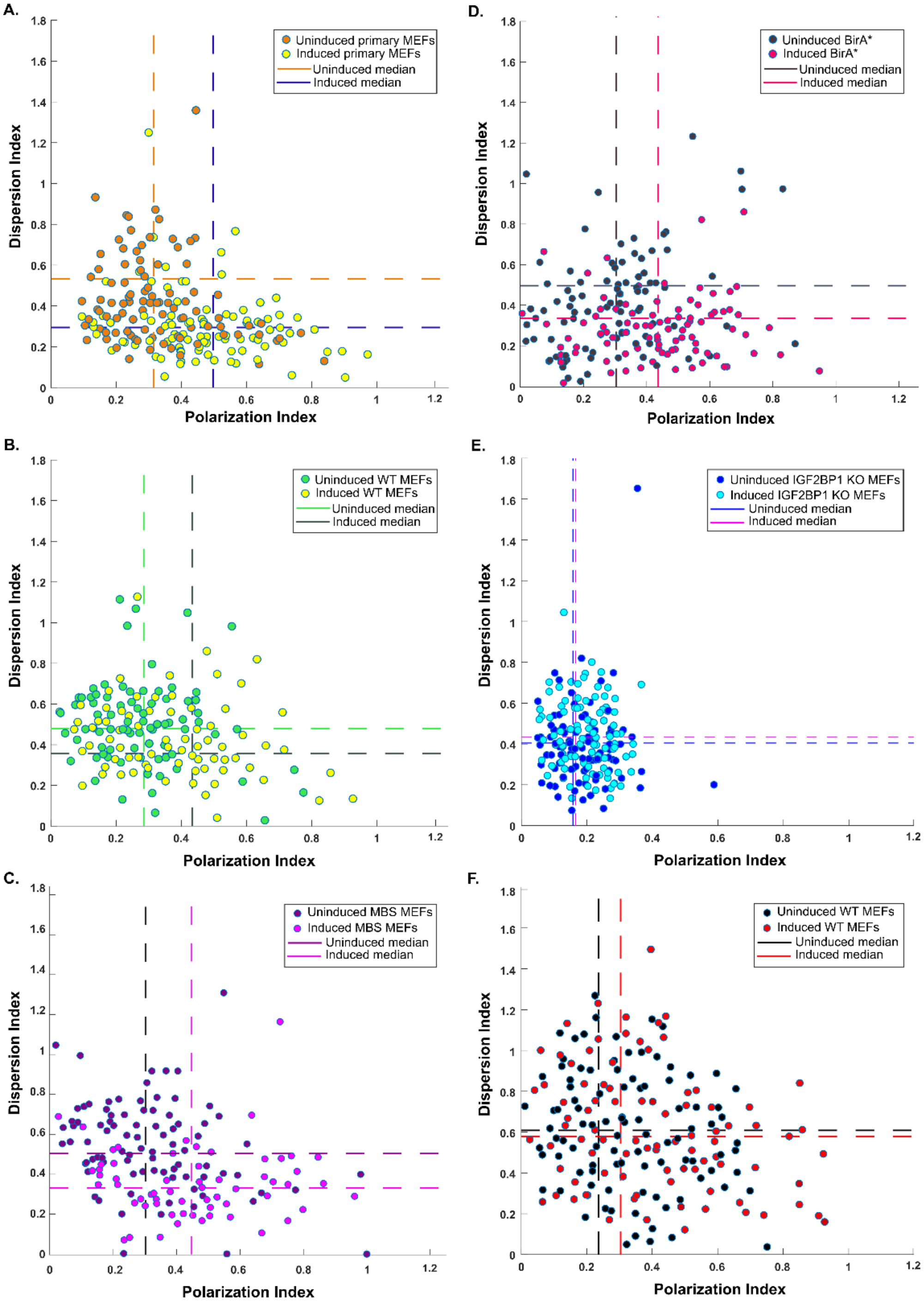


**Fig.S1**

**Scatter plot to compare changes in β-actin or Gapdh mRNA distribution.** Scatter plot displays polarization index for β-actin mRNA (A-E) or Gapdh (F) mRNA on X axis against dispersion index on y axis. Data in A - E are derived from smFISH images detecting β-actin mRNA in primary MEFs, immortalized (WT) MEFs, MBS MEFs, MBS-BirA* MEFs, or IGF2BP1 knockout MEFs. Data in F are derived from smFISH images detecting Gapdh mRNA in immortalized (WT) MEFs. 100 cells were counted for each growth condition (serum induced or uninduced).

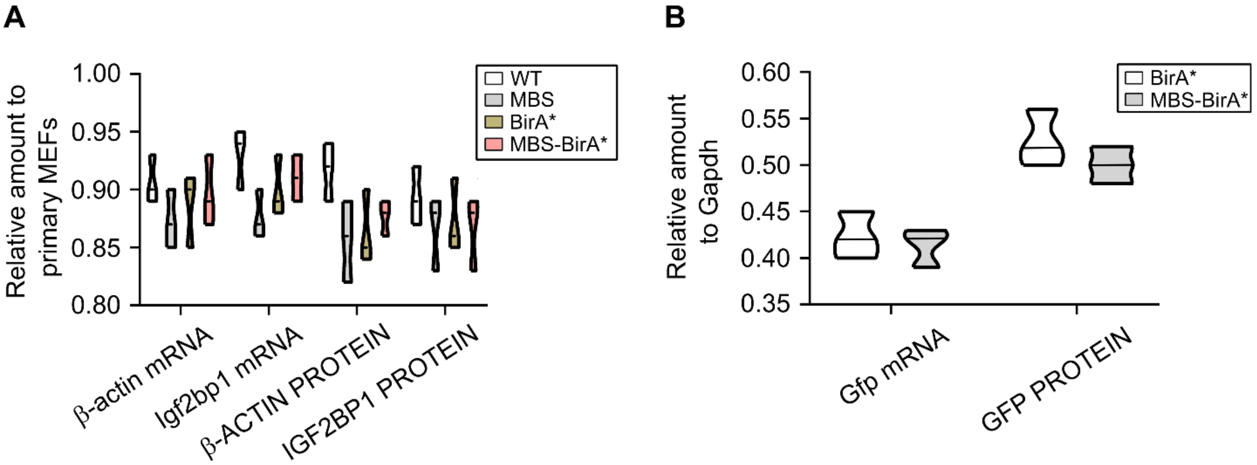


**Fig.S2**

**Violin plot to compare changes in β-actin or Igf2bp1 or heterogenous Gfp mRNA and protein expression in various cell types.**

(A) Violin plot displaying amounts of endogenous β-actin or Igf2bp1 mRNA and protein (compared to primary MEFs). (B) Violin plot displaying amounts of heterogenous GFP mRNA (normalized to endogenous Gapdh mRNA or protein levels) Data relates to Fig. 1D and shows measurements in primary MEFs, immortalized (WT) MEFs, MBS MEFs, MBS-BirA* MEFs, or BirA* MEFs. The black lines represent median values derived from 3 biological replicates.

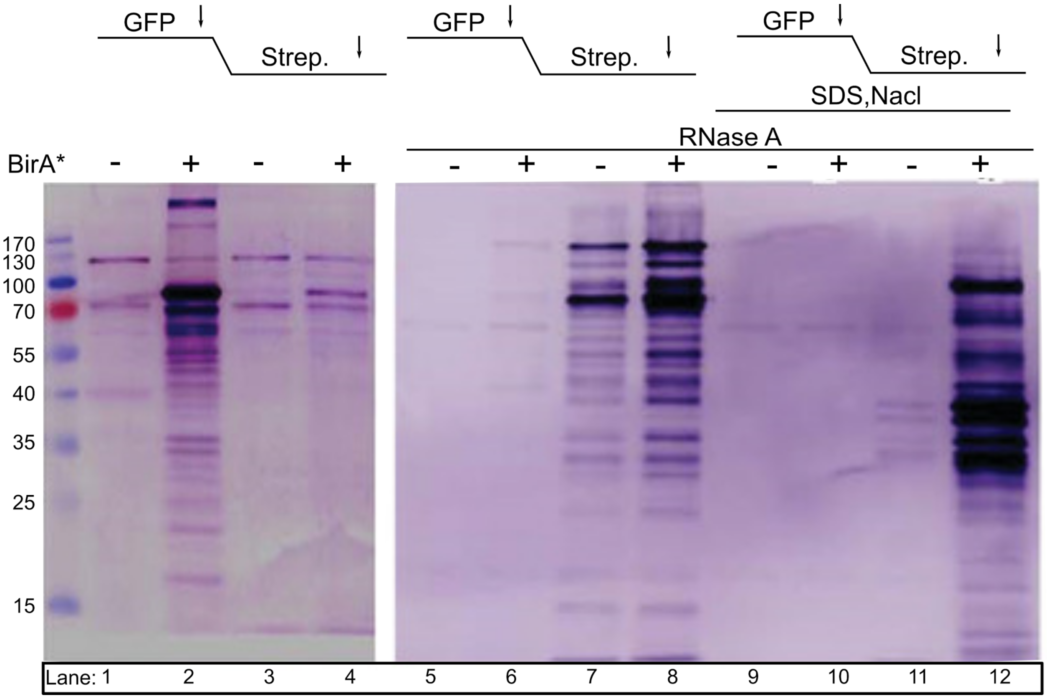


**Fig.S3**

**Optimization of purification schedule for biotinylated proteins labeled by RNA-BioID.** Specific enrichment of β-actin-MBS associated, biotinylated proteins is achieved by stringent conditions during purification. Two consecutive affinity purifications (anti-GFP followed by streptavidin pulldown) were performed. Western blots were stained for biotinylated proteins by streptavidin-alkaline peroxidase. Left panel: The majority of the biotinylated proteins remain associated with 2xMCP-eGFP-BirA* in the GFP pulldown fraction under low-stringency purification conditions (lane 2). Combination of treatment with RNase A (lane 6 versus 8), or 0.5% SDS and 500 mM NaCl (lane 10 versus 12) leads to quantitative enrichment of biotinylated proteins by streptavidin pulldown.

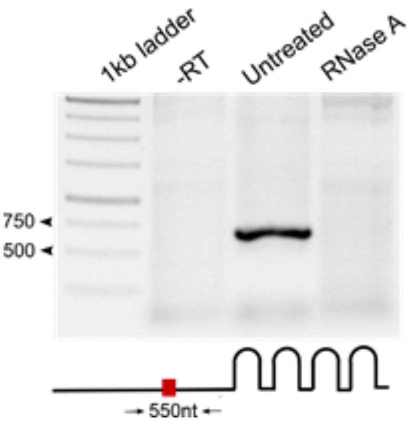


**Fig.S4**
**Validation of β-actin mRNA degradation.** β-actin-MBS-BirA* and β-actin-MBS-eGFP expressing cells were treated with 100 µg/ml RNase A for 30 min at 37°C before affinity purification (Fig. S1). Efficient RNA removal was checked by RT-PCR using a primer set that spans the indicated 550 nucleotide region upstream of the MBS cassette.

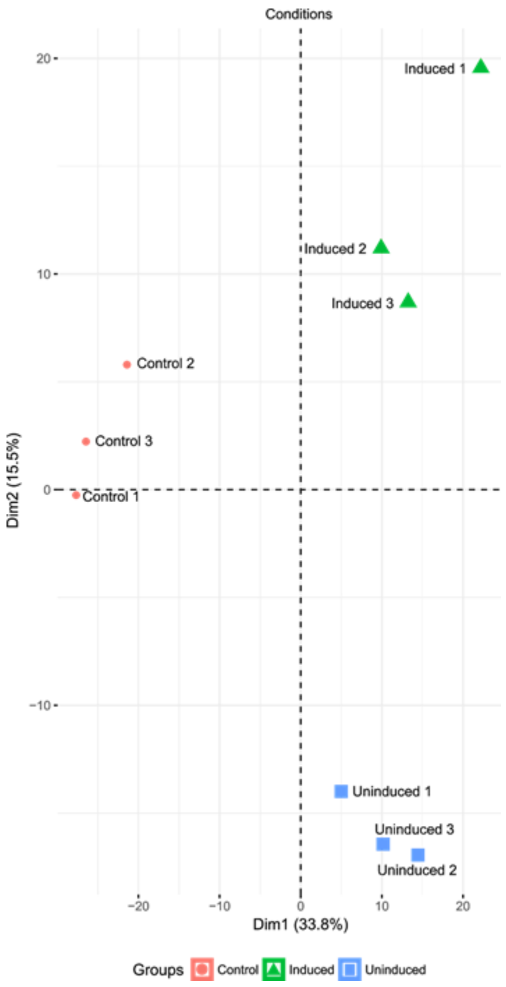


**Fig.S5**

**Visualization of sample correlation by principal component analysis (PCA).** PCA analysis reveals the largest variance between control and induced / uninduced samples on dimension 1 (variation explained 33.8%) and further separates control, uninduced and induced samples on dimension 2 (variation explained 15.5%).

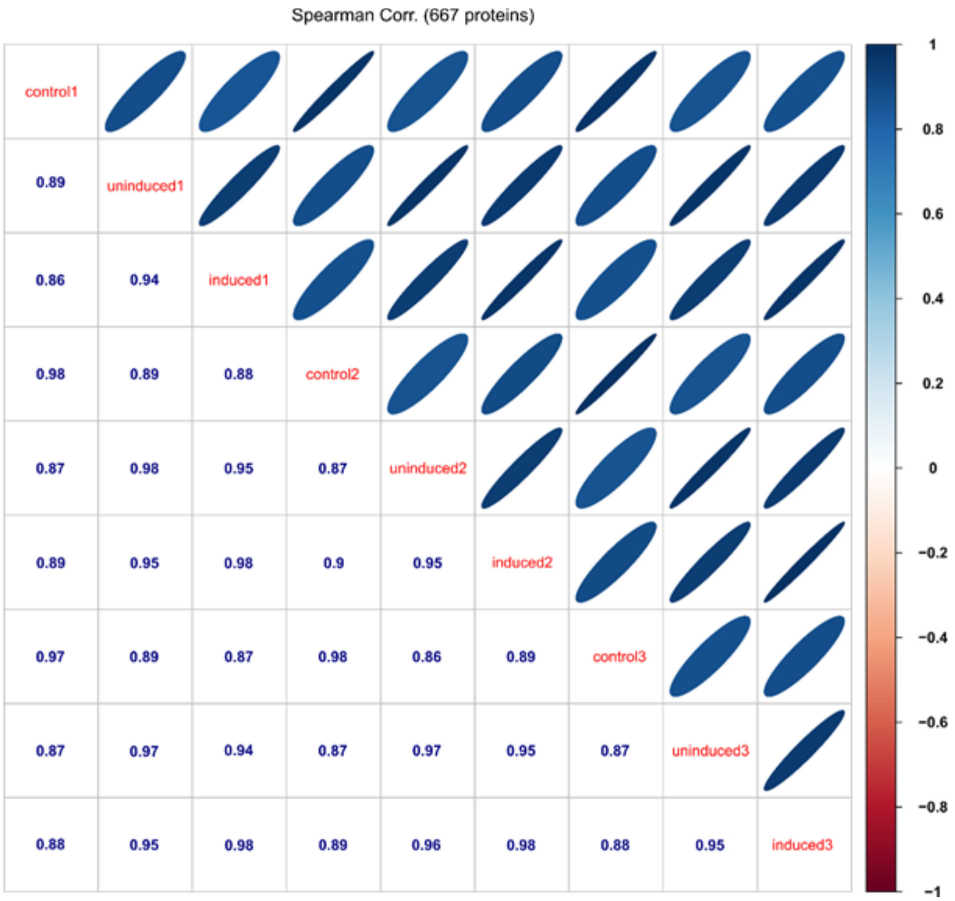


**Fig.S6**

**Spearman correlation calculation between all possible combinations of control (MCP-eGFP-BirA* only), induced and uninduced MEF samples (Control, induced, uninduced).** Correlation calculation was done based on LFQ values for common proteins. The color gradient represents high correlation (dark blue) to high anti-correlation (dark red). The analysis shows that there is a very high correlation within biological replicates per condition (>0.97), which shows the high reproducibility of the datasets.

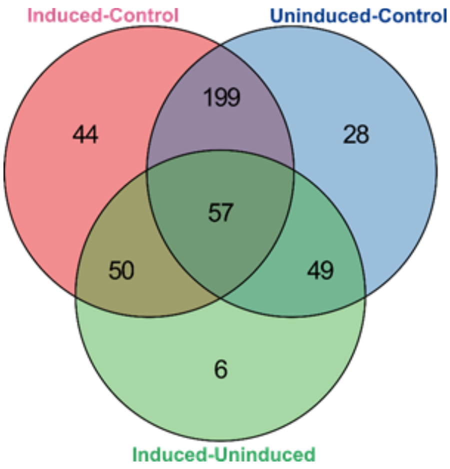


**Fig.S7**
**Venn-diagram representation of significantly enriched proteins in each population of MEFs (control/induced/uninduced).** In total, 350 proteins were significantly enriched in the dataset of induced MEFs versus control, 333 proteins in the dataset of uninduced MEFs versus control, and 162 proteins in the dataset of induced versus uninduced MEFs. 57 proteins were common between all three datasets. 44 unique proteins were identified in induced MEFs versus control, 28 proteins in uninduced MEFs versus control and six proteins were unique to the dataset of induced MEFs versus uninduced.

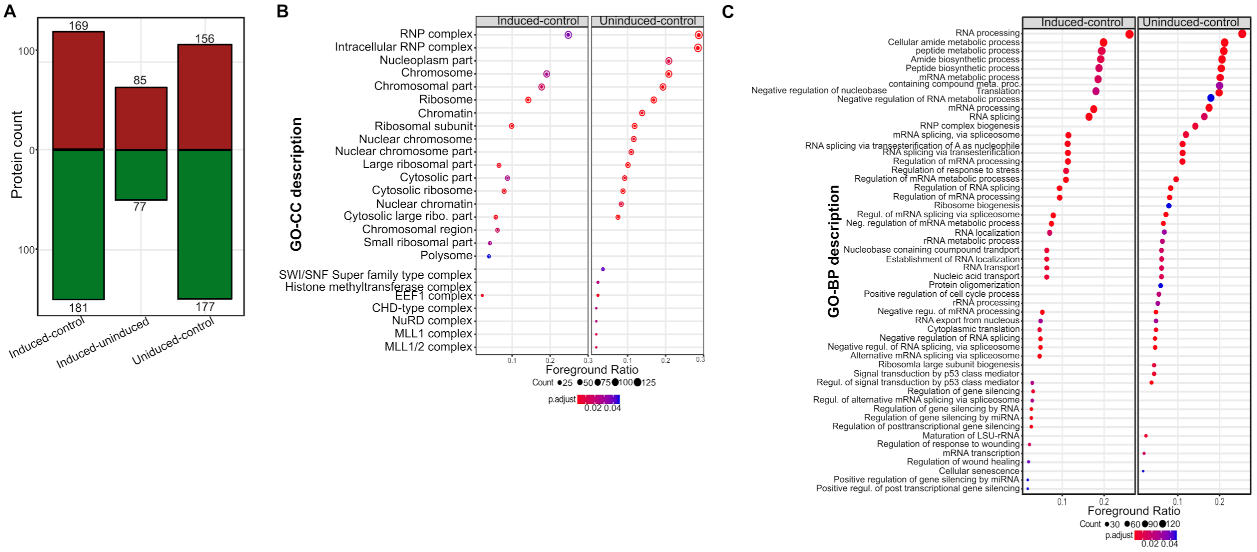


**Fig.S8**
**GO term analysis of the proteins found in MEFs expressing β-actin-24MBS and MCP-eGFP-BirA* under two different conditions (uninduced, and serum-induced MEFs) versus control MEFs.**
(A) Comparison of the number of significantly enriched proteins between all conditions. 169 proteins were significantly enriched in the induced sample (left bar) over control, while 156 proteins were overrepresented in uninduced over control (right bar). 85 proteins were enriched in the serum induced fractions over samples from uninduced MEFs (middle bar).
(B) Gene Ontology (GO) over-representation analysis of the proteins enriched in uninduced and induced conditions compared to the control. The foreground ratio represents the number of significantly enriched proteins divided by the number of proteins within each GO term. Color gradient represents the adjusted p-value (threshold adj. p-value ≤ 0.05). Ranking of gene ontology term for cellular components (GO-CC) based on foreground ratio revealed 'RNP complex" as one of the most over-represented function.
(C) Similar analysis as in (b) for gene ontology molecular function terms (GO-MF).

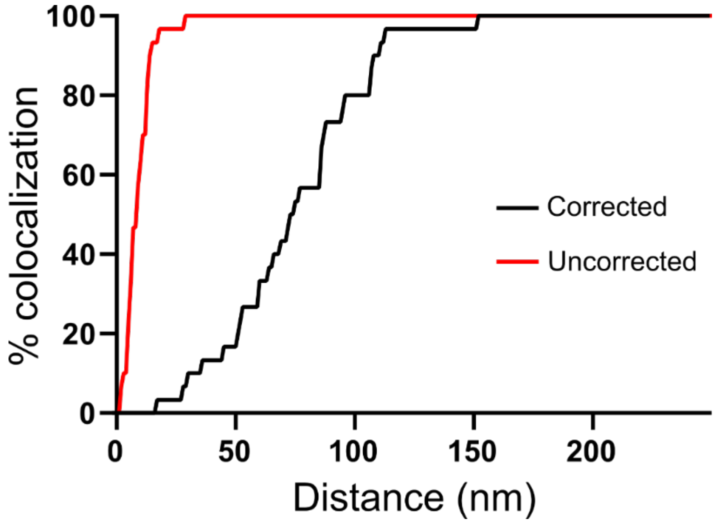


**Fig S9.**
**Correction for chromatic and mechanical shifts by microsphere fluorescent beads** Percentages of co-localization within spectrally separated centroids (collected from the beads) before (black line) and after (red line) correction was applied to the entire field of view.

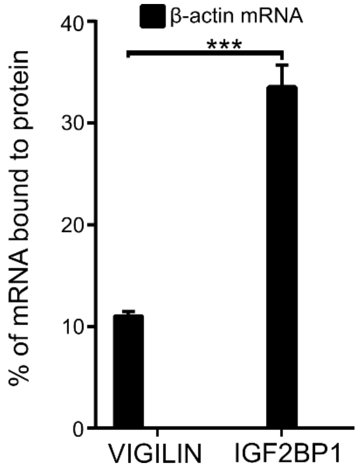


**Fig S10.**
**Co-immunoprecipitation of β-actin mRNA with IGF2BP1 and VIGILIN.**
Bars represent percentage of input mRNA co-purifying with the indicated protein. IGF2BP1 binds to around 33% of endogenous β-actin mRNA while VIGILIN was associated with 10% of endogenous β-actin mRNA which is non-significant compared to the binding efficiency of IGF2BP1 with the same mRNA. Error bars represents mean ± sem from three independent experiments. Statistical significance of each dataset was determined by Student's t-test; ***$P < 0.001$

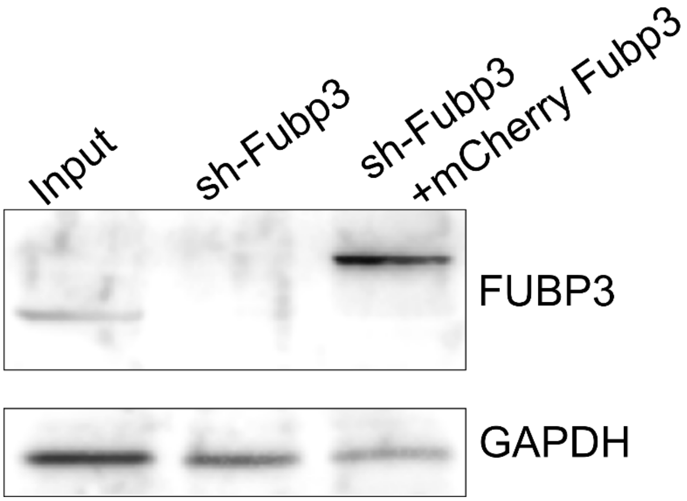


**Fig S11.**
**FUBP3 expression in rescue experiment (related to Fig.6).** Compared to input (lane 1: untreated), expression of FUBP3 or mCherry-FUBP3 was checked by western blot in MEFs with a stably transfected shFubp3 knockdown construct (lane 2) or in MEFs carrying this construct and overexpressing a knockdown-resistant mCherry-FUBP3.

Fig.1D

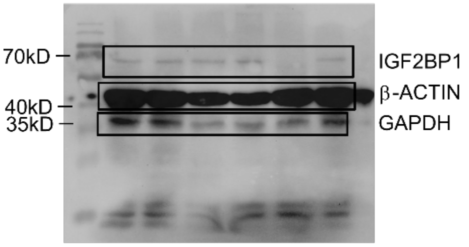


Fig.4B

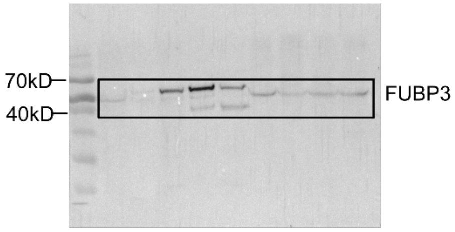


Fig.4B

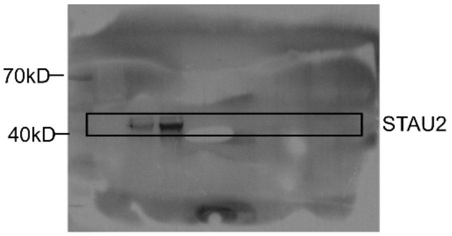


Fig.5C

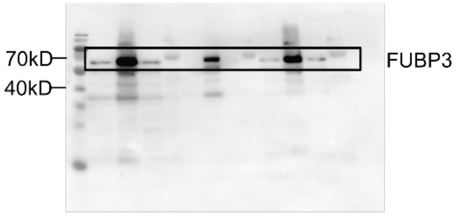


Fig.5C

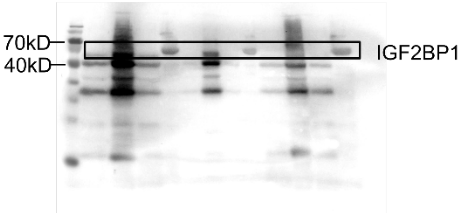


Fig.1G

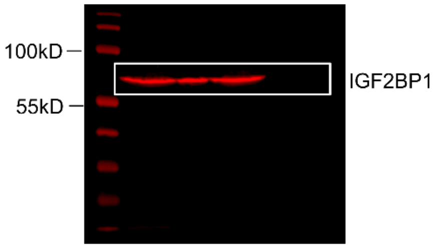


Fig.6A

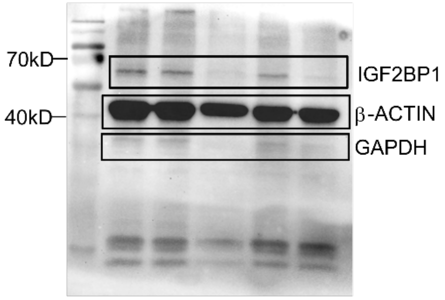


Fig. S10

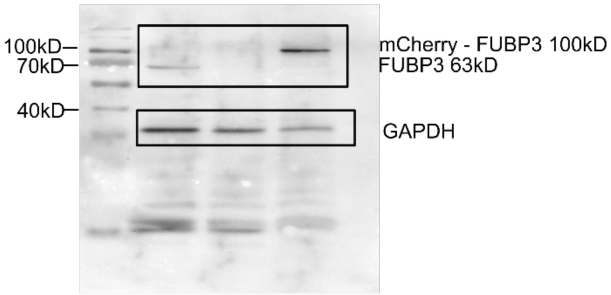


**Fig S12.**
**Uncropped western blots related to indicated figures in the manuscript.**

**Supplementary Tables**

**Supplementary table 1: Primers and oligos used to generate plasmids**

| Plasmid | Forward primers/oligos | Reverse primers /oligos |
|---|---|---|
| pHAGE-NLS-2XMCP-eGFP | GCCTCTAGAATGGTGAGCAAGGGCGAGGAGCTGTT | CCGGATCGATTTACTTGTACAGCTCGTCCATG |
| pHAGE-NLS-2XMCP-eGFP-BirA* | GCCTCTAGAATGGTGAGCAAGGGCGAGGAGCTGTT | CCGATCGATTTACTTCTCTGCGCTTCTCAGGGAGATTT |
| N-terminal mCherry pcDNA3.1(+) | CTAGCTAGCATGGTGAGCAAGGGCGAGGAGGATAAC | CCCCTTAAGCTTGTACAGCTCGTCCATGCCGCCGG |
| mCherry Fubp3 pcDNA3.1(+) | CGGGGTACCATGGCGGAGCTGGTGCAGGGGCAGAGCGCTCCGGTGGG | ATAAGAATGCGGCCGCCTGCTCCTGGCTGTGGGCCTGAGCCTGCCC |
| mCherry Fubp3 KH1mt pcDNA3.1(+) | GTTGGATTTATTATTGGCGACGACGGTGAGCAGATTTCACGA | TCGTGAAATCTGCTCACCGTCGTCGCCAATAATAAATCCAAC |
| mCherry Fubp3 KH2 pcDNA3.1(+) | GTGGGGCTGGTCATCGGCGACGACGGGGAAACGATCAAGCAG | CTGCTTGATCGTTTCCCCGTCGTCGCCGATGACCAGCCCCAC |
| mCherry Fubp3 KH3 pcDNA3.1(+) | GTTGGGATTGTCATAGGAGACGACGGAGAAATGATTAAGAAG | CTTCTTAATCATTTCTCCGTCGTCTCCTATGACAATCCCAA |
| mCherry Fubp3 KH4 pcDNA3.1(+) | GTGCGGCCTTGTGATAGGCGACGACGGCGAGAACATCAAAAGCA | TGCTTTTGATGTTCTCGCCGTCGTCGCCTATCACAAGGCCGCAC |
| mCherry Fubp3 4KH mt pcDNA3.1(+) | CGGGGTACCATGGCGGAGCTGGTGCAGGGGCAGAGCGCTCCGGTGGG | ATAAGAATGCGGCCGCCTGCTCCTGGCTGTGGGCCTGAGCCTGCCC |
| Mod-pcDNA3.1(+) | CGGGCTAGCGTTTAAACTTAAGCTTGGTACCGAGCTCGG | CCATCGATATTTCGATAAGCCAGTAAGCAGTGGGTTCTC |
| β-actin-3’UTR mod-pcDNA3.1(+) | CCATCGATTAATACGACTCACTATAGGGCGGACTGTTACTGAGCTGCG | CGGGCTAGCTGTTTGTGTAAGGTAAGGTGTGCACTTTTATTG |
| Fubp3 pETM14 | CATGCCATGGATGGCGGAGCTGGTGCAGGGGC | ATAAGAATGCGGCCGCCTGCTCCTGGCTGTGGGCCTGAGCCTGCCC |

| | | |
|---|---|---|
| Igf2bp1 pETM14 | CATGCCATGGACAAGCTTTACATCGGCAACCTCAACGAGAGTG | GCCGAATTCCTTCCTCCGAGCCTGGGCCAGGTTGCT |
| pHAGE-mCherry FUBP3 shRNA rescue | F1: AGATGCCAACACGCCGCCAGAATCATCAATGAGCTCATTCTC<br>F2: : CGGGGTACCATGGCGGAGCTGGTGCAGGGG | R1: GAGAATGAGCTCATTGATGATTCTGGCGGCGTGTTGGCATCT<br>R2: AAGGAAAAAAGCGGCCGCCTGCTCCTGGCTGTG |

**Supplementary table 2: Primers used for PCR or qRT-PCR**

| Gene/Insert | Forward primers/oligos | Purpose |
|---|---|---|
| eGFP F | GCCTCTAGAATGGTGAGCAAGGGCGAGGAGCTGTT | PCR |
| eGFP R | GGCCTCGAGCTTGTACAGCTCGTCCATGCCGAGAGT | PCR |
| BirA* F | GCCCTCGAGGACAAGGACAACACCGTGCCCCTGAA | PCR |
| BirA* R | CCGATCGATTTACTTCTCTGCGCTTCTCAGGGAGATTT | PCR |
| Igf2bp1 F | TACAAGTGTTCATCCCCGCC | q-RT |
| Igf2bp1 R | AGGTGTTTCTGGTGGTGCAA | q-RT |
| β-actin R | GCCTTCACCGTTCCAGTTTTT | q-RT |
| β-actin F | CTGAGCTGCGTTTTACACCC | q-RT |
| β-actin F | CTGAGAGGGAAATCGTGCGT | RT |
| β-actin R | AGGGTGTAAAACGCAGCTCAG | RT |
| Gapdh F | GAGGGATGCTGCCCTTACC | q-RT |
| Gapdh R | AAATCCGTTCACACCGACCT | q-RT |
| Fubp3 F | TAGTACTCAGCCCAGGCCAT | q-RT |
| Fubp3 R | GTAGTGACTGGAGTGTGGGC | q-RT |
| β-actin zip code F | CGGACTGTTACTGAGCTG | q-RT |
| β-actin zip code R | CGCAAGTTAGGTTTTGTC | q-RT |
| β-actin proximal zip code F | GCAGAAAAAAAAAAAAT | q-RT |

| | | |
|---|---|---|
| β-actin proximal zip code R | ACAAAGCCATGCC | q-RT |
| β-actin mutant zip code F | GTTACTGAGCTGCGT | q-RT |
| β-actin mutant zip code R | CGCAAGTTAGGTTTTGTC | q-RT |
| β-actin 3' UTR F | CGGACTGTTACTGAGCTG | q-RT |
| β-actin 3' UTR R | GCCTTCACCGTTCCAGTTTTT | q-RT |
| Sense-T7 Fubp3 bind-F | CGGCTAATACGACTCACTATAGGCCCCGGGGAAGGTGACAGCATTGCTTCTGTGTAAATTATGTACTGCAAAAATTTTTTTAAATCTTCCGCCTTAATAC | In vitro transcription |
| Sense-Fubp3 bind T7-R | GTATTAAGGCGGAAGATTTAAAAAAATTTTTGCAGTACATAATTTACACAGAAGCAATGCTGTCACCTTCCCCGGGGCCTATAGTGAGTCGTATTAGCCG | In vitro transcription |
| Sense-T7 Fubp3 mt bind-F | CGGCTAATACGACTCACTATAGGCCCCGGGGAAGGTGACAGCATTGCTTCTGTGTAAATACTGCAAAAATTTTTTTAAATCTTCCGCCTTAATAC | In vitro transcription |
| Sense-Fubp3 mt bind T7-R | GTATTAAGGCGGAAGATTTAAAAAAATTTTTGCAGTATTTACACAGAAGCAATGCTGTCACCTTCCCCGGGGCCTATAGTGAGTCGTATTAGCCG | In vitro transcription |
| T7-Zipcode-F | CCTAATACGACTCACTATAGGCGGACTGTTACTGAGCTGCGTTTTACACCCTTTCTTTGACAAAACCTAACTTGC | In vitro transcription |
| T7-Zipcode-R | GCAAGTTAGGTTTTGTCAAAGAAAGGGTGTAAAACGCAGCTCAGTAACAGTCCGCCTATAGTGAGTCGTATTAGG | In vitro transcription |
| T7-Zipmt-F | CCTAATACGACTCACTATAGGGTTACTGAGCTGCGTTTTTTTCTTTGACAAAACCTAACTTGC | In vitro transcription |
| T7-Zipmt-R | GCAAGTTAGGTTTTGTCAAAGAAAAAAACGCAGCTCAGTAACCCTATAGTGAGTCGTATTAGG | In vitro transcription |
| T7-zipcode proximal-F | CCTAATACGACTCACTATAGGGCAGAAAAAAAAAAAATAAGAGACAACATTGGCATGGCTTTGT | In vitro transcription |
| T7-zipcode proximal-R | ACAAAGCCATGCCAATGTTGTCTCTTATTTTTTTTTTTTCTGCCCTATAGTGAGTCGTATTAGG | In vitro transcription |

**Supplementary table 3: Antibodies used in this study**

| Antibody | Host | Dilution | Buffer (for Western blot) | Supplier |
|---|---|---|---|---|
| anti-β-ACTIN | Mouse-monoclonal | 1:4000 - western | 0.3 % BSA/TBST | Clone AC15 [Sigma-Aldrich (A1978)] |
| anti-GAPDH | Mouse-monoclonal | 1:1000 - western | 0.3 % BSA/TBST | Proteintech. 60004-1-Ig. Clone no. 1E6D9 |
| anti-IGF2BP1 | Mouse | 1:4000 - Western<br>1:2000 - IF | 0.3 % BSA/TBST | [MBL (RN001M)] |
| anti-FUBP3 | Rabbit monoclonal | 1:4000 - Western<br>1:1000 - IF | 0.3 % BSA/TBST | Abcam: ab181122 |
| anti-GFP | Chicken | 1:5000 - IF | | Aves Labs, Inc. (GFP-1010)] |
| anti-GFP | Rabbit polyclonal | 1:4000 - Western | 0.3 % BSA/TBST | Invitrogen- A-6455 |
| anti-AP streptavidin | | 1:4000 | 100 mM NaCl, 50 mM MgCl2, 100 mM Tris-Cl pH 9.5 | |
| anti-Mouse IgG HRP | Sheep | 1:10.000 | 1 % BSA/TBST | Jackson-ImmunoResearch: 515-035-003 |
| anti-Rabbit IgG HRP | Goat | 1:10.000 | 1 % BSA/TBST | Jackson-ImmunoResearch:111-055-144 |
| Alexa Fluor-labeled anti-chicken, -mouse, or -rabbit secondary antibodies | Goat | 1:1000 - IF | | Life Technologies |

IF: Immunofluorescence

**Supplementary table 4: Probes used for smFISH**

smFISH probes used to detect the MS2 binding site (labeled on both 5' and 3' with ATTO488)

| | |
|---|---|
| MS2_LK20 | TTTCTAGAGTCGACCTGCAG |
| MS2_LK51_1 | CTAGGCAATTAGGTACCTTAG |
| MS2_LK51-2 | CTAATGAACCCGGGAATACTG |

smFISH probes used to detect the β-actin ORF (labeled on 3' with ATTO633)

| | |
|---|---|
| Actb807 | ATAGTGATGACCTGGCCGTCAG |
| Actb830 | CATCGGAACCGCTCGTTGCC |
| Actb863 | ACCCAAGAAGGAAGGCTGGAA |
| Actb886 | TCATGGATGCCACAGGATTCC |
| Actb927 | CGGATGTCAACGTCACACTTCA |
| Actb960 | CCAGACAGCACTGTGTTGGCAT |
| Actb984 | ATGCCTGGGTACATGGTGGTAC |
| Actb1007 | TCTCCTTCTGCATCCTGTCAGC |
| Actb1034 | CATGGTGCTAGGAGCCAGAGC |
| Actb1073 | CAGAGTACTTGCGCTCAGGAGG |
| Actb1099 | CCAGGATGGAGCCACCGATC |
| Actb1122 | ATCTGCTGGAAGGTGGACAGTG |
| Actb1145 | CGTACTCCTGCTTGCTGATCCA |
| Actb1178 | CTTGCGGTGCACGATGGAGGG |
| Actb1208 | CGCAGCTCAGTAACAGTCCGC |
| ACTB1228 | TCAAAGAAAGGGTGTAAAAC |
| ACTB1254 | TTTTTTTTTTTTCTGCGCAAGTTAG |
| ACTB1284 | AAAGCCATGCCAATGTTGTC |
| ACTB1355 | GCGCCAAAACAAAACAAAAAAACTTA |
| ACTB1394 | TCACCGTTCCAGTTTTTAAA |

| | |
|---|---|
| ACTB1423 | ATGTTTGCTCCAACCAACTG |
| ACTB1446 | CCACATTTGTAGAACTTTGG |
| ACTB1470 | CAAAACAATGTACAAAGTCC |
| ACTB1511 | GGAATGACTATTAAAAAAAGAC |
| ACTB1535 | ACCACTTATTTCATGGATAC |
| ACTB1577 | AGGAGTGGGGGTGGCTTTTG |
| ACTB1598 | GGACGCGACCATCCTCCTCT |
| ACTB1626 | ACCTTCCCCGGGGTGGACT |
| ACTB1661 | ATTTTTGCAGTACATAATTTACAC |
| ACTB1683 | TTAAGGCGGAAGATTTAAAAAAA |
| ACTB1724 | ACCTGGGCCATTCAGAAATT |
| ACTB1749 | GGGACAAAAAAAAGGGAGGC |
| ACTB1787 | CTCCCAGGGAGACCAAAGCC |
| ACTB1810 | GGCTGCCTCAACACCTCAAC |
| ACTB1831 | GTCAGTGTACAGGCCAGCCC |
| ACTB1855 | GGTGTGCACTTTTATTGGTC |

### Supplementary Methods

**Plasmids and cloning:** The lentivirus vector pHAGE-UbiC carrying NLS-2X MCP-tagRFPT (1) was used as backbone to generate all lentiviral vectors used in this study. In order to generate pHAGE-NLS-2XMCP-GFP, the GFP fragment was amplified from pEGFPC1 and cloned into the lentiviral vector within XbaI and ClaI restriction sites, thereby replacing the tagRFPT sequence. To generate pHAGE-NLS-2XMCP-eGFP-BirA*, a BirA* fragment was amplified from plasmid pSF3-TGN38-cMyc-BirA* (2) with XhoI and ClaI sites, eGFP fragment was amplified with XbaI and XhoI sites, after digestion both the fragments were ligated and the ligated product was PCR amplified using forward XbaI eGFP primer and reverse ClaI BirA* primer and the amplified product was integrated into pHAGE-NLS-2XMCP-eGFP plasmid under XbaI and ClaI sites.
Plasmids expressing mCherry fusion proteins were generated on pcDNA3.1(+) backbone. The mCherry CDS was amplified from plasmid pmCherry-C1 (Clontech), introducing NotI and XhoI sites at the 5' or 3' ends and the PCR product ligated into pcDNA3.1(+) to generate a C-terminal tag mCherry pcDNA3.1(+) entry plasmid. To create a N-terminal tag mCherry pcDNA3.1(+) entry plasmid, mCherry was amplified with NheI-mCherry forward and AflIII-mCherry reverse primer and cloned at the beginning of the MCS of pcDNA3.1(+).
Coding sequences of Igf2bp1 (Acc.No. NM_009951.4), and Fubp3 (Acc.No. NM_001290548.1), were amplified from Mouse (C57BL/6J) cDNA and cloned into C-terminal mCherry using restriction sites NheI and EcoRI (for Igf2bp1), KpnI and NotI (Fubp3).
To create plasmids expressing KH mutants of Fubp3-mCherry, Fubp3-mCherry was chosen as the template and different primer sets (Supplementary table 2) were used for inverse PCR to change the G-X-X-G motifs to G-D-D-G (3). After treating the PCR reaction with DpnI for overnight at 37°C, the linear amplification product was transformed into E.coli Dh5α for re-ligation and amplification.
To create a backbone vector for in vitro transcription, plasmid pcDNA3.1 was modified (mod-pcDNA3.1(+)) to remove unwanted restriction sites and to introduce a new restriction site before the T7 promoter. Two primers were designed for inverse PCR of pcDNA3.1(+), resulting in an additional ClaI site in front of the T7 promoter while at the same time removing a part of the multiple cloning site. DNA sequences encoding the β-actin zipcode, the proximal zipcode or a mutant zipcode were generated by annealing two complementary oligos containing NheI-ClaI overhangs at the ends and after phosphorylation ligated into the mod-pcDNA3.1(+) vector within NheI and ClaI sites. To generate pcDNA3.1(+) β-actin-3'UTR, the complete β-actin 3'UTR was amplified and cloned into the same sites.

**Cell culture and serum starvation assay:** Mouse embryonic fibroblasts (MEFs) from C57BL/6J mice or Hek293T cells were cultured in DMEM (with 4.5 g glucose and L-glutamine containing 10% fetal bovine serum (FBS) and 1% pen-strep). For serum starvation of MEFs, cells were starved in DMEM medium without serum (including 1% pen-strep) for at least 24 hrs and induced with DMEM media containing 10% FBS and 1% pen- strep for different time points as mentioned in the results section.

**Lentivirus generation:** Lentiviral particles were produced by transfecting the expression vector along with plasmids (Addgene plasmid numbers 12259, 12251, and 12253) for ENV (pMD2.VSVG), packaging (pMDLg/pRRE), and REV (pRSV-Rev) into HEK293T cells using Fugene HD reagent (Promega). The virus-containing supernatant was harvested on day 2 and day 3 and centrifuged at 500 x g for 10 min at 4˚ C and passed through 0.45 µm filter. The filtered supernatant was used for transduction of MEFs cells at a dilution of 1:100 and in presence of 8 mg/ml protamine sulphate. 3 days after the transfection, MEFs were washed 3x with serum containing media, trypsinized and washed twice again. Cells were washed once in FACS buffer (1x DPBS without calcium and magnesium, 0.2% BSA, 0.5 mM EDTA, 5 mM $MgCl_2$) and resuspended in FACS Cells expressing low levels of GFP were isolated with a FACS Aria cell sorter (Becton-Dickinson) before further culturing.

shRNA knockdown of Fubp3 and generation of FUBP3 rescue cell line. For generating stable cell lines expressing an shRNA construct against FUBP3 (Dharmacon clone ID. V3IMMMCG_11752124), GAPDH (Dharmacon clone ID. VSM11618), or a non-targeting control (Dharmacon clone ID. VSM11618), SMARTvector™ plasmids expressing inducible lentiviral shRNAs and shMIMIC™ inducible lentiviral microRNA were stably integrated in WT MEFs. For selection of positive cell lines 2μg/ml concentration of puromycin was chosen to establish stably integrated cell lines. For induction, dose and time response curves for doxycycline were generated and an incubation for three days with a doxycycline concentration of 200 ng/ml was chosen for the knockdown.
For the FUBP3 rescue cell line, a plasmid was generated using the PHAGE-UBC lentiviral backbone. The rescue construct contained an N-terminal mCherry sequence and silent mutations (CGG TGT CAG CAT GCA GCT CGC) in the shRNA target sequence (CGG TGC CAA CAC GCC GCC AGA). The target region is starting at base pair 926 n of the Fubp3 CDS. The resulting plasmid was stably integrated in FUBP3-shRNA MEFs and positive cells were collected by FACS sorting against mCherry.

**Immunoprecipitation, Western blot, and qRT-PCR:** For immunoprecipitation, cells were lysed in IP lysis buffer polysome extraction buffer (PEB) (without cycloheximide), atleast 200 μg of total protein was used per pull down experiment. For the rest of the experiment including western and qPCR, the protocol was followed as mentioned before (4) with the following modifications. A total of 200 μg of proteins were taken after lysing the cells in IP lysis buffer. 100 μl of protein A (for anti-mouse antibodies) or protein G (for anti-rabbit antibodies) coupled magnetic beads were used for IP. After washing them 3x in NT2 buffer (4) the beads were blocked with 5% BSA and 0.5 mg/ml ssDNA. After washing the beads once again with NT2 buffer, the beads were incubated with either 20 μg of FUBP3 antibody or 10 μg of IGF2BP1 in NT2 buffer at a total volume of 200μl, for overnight at 4°C with end to end rotation. After preclearing the lysates with 100μl of protein A or Protein G mag beads, antibody coupled beads were added in the lysates and incubated for 4 hrs at 4°C with end to end rotation. After the incubation, beads were separated from the lysates by a magnetic stand and washed 5 times with ice cold NT2 buffer and the supernatant was removed. Beads were resuspended in 100 μl of NT2 buffer. For isolation of proteins from the beads, 40 μl of beads were boiled in 100 μl 1x Laemmli buffer for 10 min at 95 C and the elute was separated from the beads on a magnetic strand. For western blots 40 μl from the eluted sample were used. For RNA isolation, in the 60 μl of the remaining beads, 5 μl proteinase K (10 mg/ml) and 1 μl of 10% SDS were added and incubated at 55°C water bath for 30 min. 100 μl of buffer NT2 was added in the sample and 200 μl of acidic phenol-chloroform mix was added and vortexed for 10 sec. After centrifugation at 16,000g at room temperature for 5 min, the upper aqueous layer was isolated, sodium acetate (pH 5.2, final concentration 0.3 M), 5 μl glycoblue (Invitrogen), and 600 μl of ethanol was added to precipitate the RNA. The RNA pellet was resuspended in 50 μl of RNase free water. For qPCR analysis, 500 ng of RNA was used after treating with 10U of DNase I.

**NanoLC-MS/MS analysis and data processing:** Beads were resuspended in denaturation buffer (6 M urea, 2 M thiourea, 10 mM Tris buffer, pH 8.0), and proteins were reduced by incubation in 1 mM dithiothreitol (DTT) for 1 h at room temperature. Alkylation of reduced cysteines was performed in 5.5 mM iodoacetamide (IAA) in 50 mM ammonium bicarbonate (ABC) buffer for 1 h at room temperature in the dark. On beads digestion of proteins was started with endoproteinase LysC (2 μg per 100 μg protein) for 3 h of incubation at pH 8.0 and room temperature. Tryptic digestion (2 μg per 100 μg protein) was performed overnight at room temperature after diluting the sample with four volumes of 20 mM ABC and adjusting the pH to 8.0. Acidified peptides were purified via PHOENIX Peptide Clean-up Kit (PreOmics) according to user manual, and separated on an EasyLC nano-HPLC (Thermo Scientific) coupled to an LTQ Orbitrap Elite (Thermo Scientific) as described elsewhere (5) with slight modifications: The peptide mixtures were injected onto the column in HPLC solvent A (0.1% formic acid) at a flow rate of 500 nl/min and subsequently eluted

with an 87 minutes segmented gradient of 5–33-50-90% of HPLC solvent B (80% acetonitrile in 0.1% formic acid) at a flow rate of 200 nl/min.
Precursor ions were acquired in the mass range from m/z 300 to 2000 in the Orbitrap mass analyzer at a resolution of 120,000. Accumulation target value of 106 charges was set. The 15 most intense ions were sequentially isolated and fragmented in the linear ion trap using collision-induced dissociation (CID) at the ion accumulation target value of 5000 and default CID settings. Sequenced precursor masses were excluded from further selection for 60 s. Acquired MS spectra were processed with MaxQuant software package version 1.5.2.861 with integrated Andromeda search engine (6). Database search was performed against a target-decoy Mus musculus database obtained from Uniprot, containing 60,752 protein entries, the sequence of MCP-eGFP-BirA and 284 commonly observed contaminants. Endoprotease trypsin was defined as protease with a maximum of two missed cleavages. Oxidation of methionine and N-terminal acetylation were specified as variable modifications, whereas carbamidomethylation on cysteine was set as fixed modification. Initial maximum allowed mass tolerance was set to 4.5 ppm (for the survey scan) and 0.5 Da for CID fragment ions. Peptide, protein and modification site identifications were reported at a false discovery rate (FDR) of 0.01, estimated by the target/decoy approach (7). The label-free algorithm was enabled, as was the “match between runs” option (8).

**Downstream analysis and functional interpretation:** Downstream analyses were performed in the R environment. The resultant proteome profiles obtained were quality checked for replicate correlation using principal component analysis and hierarchical clustering. The data was then filtered for low abundant proteins via a two-step process. Firstly, all proteins were ranked (in descending order) by LFQ intensity and the 3rd quartile LFQ value set as the minimum intensity threshold. Secondly, to pass the filtering, all samples within a condition must have an LFQ intensity above this threshold. Contaminants and reverse hits, as well as proteins only identified by sites, were also removed from downstream analyses. Following low abundance filtering the LFQ values were quantile normalized using the MSnbase package (10). In order to statistically compare protein abundances across conditions (even when the protein was not detected in one of the conditions) imputation was used on the data using a mixed model of nearest neighbor averaging and left-censored missing data from a truncated distribution (10). Comparisons in which both conditions contained imputed values were completely discounted. Proteins which were significantly different between conditions were identified using ANOVA followed by the Tukey post-hoc test. Significance was set at an adjusted p-value of 0.05 following Benjamini-Hochberg multiple correction testing. Functional information, namely gene ontology (GO) and KEGG ID, for Mus musculus proteins were retrieved using the UniProt.ws package (11). The over-representation testing for GO and KEGG pathways were done for each comparison via the cluster Profiler package67 based on hypergeometric distribution (p-adj. ≤ 0.05). Further analysis were performed using the Perseus software (11, 12).

**Expression of recombinant proteins and *in vitro* binding assay with *in vitro* transcribed RNA:** Full length Igf2bp1, Fubp3, were cloned into vector pETM 14 containing a HIS tag. For pETM 14-Fubp3, a 1704 bp long fragment of Fubp3 was cloned into NcoI and NotI. For pETM 41-Igf2bp1, a 1724 bp long fragment of Igf2bp1 was cloned into NcoI and EcoRI sites. The proteins were expressed in Rossetta-gami™ 2(DE3) cells according to manufacturer’s protocol (Novagen). To generate constructs for *in vitro* transcription, the 683 bp long full length β-actin 3’UTR was cloned between NheI and ClaI sites into a modified pCDNA3.1(+) (described above in Plasmids and Cloning) that contains a T7 instead of the CMV promoter. For the 54 nt long localization element of β-actin, the 49 bp long region proximal to the zipcode, the mutated zipcode (13), the 79 nucleotide long region containing UAUG motif, and the 75 nucleotide long region lacking the UAUG regions, two oligonucleotides containing the corresponding sequences and a T7 promoter on the forward strand were annealed. For *in vitro* transcription, either 1 µg of the linearized plasmid (digested with NheI), or the annealed oligonucleotides were used. Transcription was done for 1 h at 37°C in 10 µl of 40 mM Tris-HCl, pH 7.5, 6 mM Mg-Acetate, 10 mM DTT, 1 mM spermidine, 0.5 mM each of ATP, GTP and

CTP, 10 µM UTP, 100 units/ml RNase inhibitor, and 500 units/ml of T7 RNA polymerase. RNA was recovered by ethanol precipitation at -20°.
For the *in vitro* binding assay, 40 µg of total protein lysates were pre-cleared with empty magnetic beads before incubation with 50 pmol of in vitro transcribed RNAs at 4°C for at least 4 hrs with end to end rotation. His-tagged proteins and bound RNAs were captured with 30µl of His60 Ni Magnetic Beads (Takara, cat no. 635693), washed twice with 1xPBS, once with buffer containing 250 mM NaCl and 10 mM Tris-HCl pH7.5 and finally once with 10 mM Tris-HCl pH7.5. Half of the beads were used for protein isolation and western blot, the rest for RNA isolation. Bound RNA was determined using qRT PCR and primers according to supplementary table 1 and 2.
